# A chromosome-length genome assembly and annotation of blackberry (*Rubus argutus*, cv. ‘Hillquist’)

**DOI:** 10.1101/2022.04.28.489789

**Authors:** Tomáš Brůna, Rishi Aryal, Olga Dudchenko, Daniel James Sargent, Daniel Mead, Matteo Buti, Andrea Cavallini, Timo Hytönen, Javier Andrés, Melanie Pham, David Weisz, Flavia Mascagni, Gabriele Usai, Lucia Natali, Nahla Bassil, Gina E. Fernandez, Alexandre Lomsadze, Mitchell Armour, Bode Olukolu, Thomas Poorten, Caitlin Britton, Jahn Davik, Hamid Ashrafi, Erez Lieberman Aiden, Mark Borodovsky, Margaret Worthington

## Abstract

**Background:** Blackberries (*Rubus* spp.) are the fourth most economically important berry crop worldwide. Genome assemblies and annotations have been developed for *Rubus* species in subgenus *Idaeobatus*, including black raspberry (*R. occidentalis*), red raspberry (*R. idaeus*), and *R. chingii*, but very few genomic resources exist for blackberries and their relatives in subgenus *Rubus*.

**Findings:** Here we present a chromosome-length assembly and annotation of the diploid blackberry germplasm accession ‘Hillquist’ (*R. argutus*). ‘Hillquist’ is the only known source of primocane-fruiting (annual-fruiting) in tetraploid fresh-market blackberry breeding programs and is represented in the pedigree of many important cultivars worldwide. The ‘Hillquist’ assembly, generated using PacBio long reads scaffolded with Hi-C sequencing, consisted of 298 Mb, of which 270 Mb (90%) was placed on seven chromosome-length scaffolds with an average length of 38.6 Mb. Approximately 52.8% of the genome was composed of repetitive elements. The genome sequence was highly collinear with a novel maternal haplotype-resolved linkage map of the tetraploid blackberry selection A-2551TN and genome assemblies of *R. chingii* and red raspberry. A total of 38,503 protein-coding genes were predicted using the assembly and Iso-Seq and RNA-seq data, of which 72% were functionally annotated.

**Conclusions:** The utility of the ‘Hillquist’ genome has been demonstrated here by the development of the first genotyping-by-sequencing based linkage map of tetraploid blackberry and the identification of several possible candidate genes for primocane-fruiting within the previously mapped locus. This chromosome-length assembly will facilitate future studies in *Rubus* biology, genetics, and genomics and strengthen applied breeding programs.

## Data Description

### Background

Blackberries (*Rubus* spp.) are specialty fruits in the *Rosoideae* subfamily of Rosaceae, which are prized for their sweet, juicy berries that have a delicate aroma and a deep black color. The global blackberry industry has experienced rapid growth and change during the past two decades [1]; Americans spent just over $656 million on blackberries during 2020, a 17% increase over the previous year [2]. This growth has been driven by increased consumer demand, advanced production methods, year-round product availability, and new cultivars.

The *Rubus* genus likely has a North American origin and is divided into 12 subgenera [3, 4]. Other economically important crops in the genus *Rubus* include red raspberries (*Rubus idaeus*) and black raspberries (*Rubus occidentalis*), both of which are diploid species belonging to subgenus *Idaeobatus.* In contrast, blackberries belong to subgenus *Rubus* and range from diploid to 12x (2n = 2x = 14 to 2n = 12x = 84). Species belonging to subgenus *Rubus* are believed to have diverged from other subgenera, including *Idaeobatus*, *Chamaebatus*, *Cylactis*, *Dalibardastrum*, and *Malachobatus,* approximately 15-20 MYA [4]. Cultivated blackberries are not assigned a specific epithet because most cultivars have several species in their ancestry [5]. In North America, erect and semi-erect blackberries grown for fresh-market production are bred at the tetraploid (2n = 4x = 28) level and are composed mostly of species native to the Central and Eastern United States, including *R. allegheniensis, R. argutus,* and *R. trivialis.* Processing cultivars with trailing growth habit are typically bred at higher ploidy levels (primarily 2n = 6x/7x= 42/49), and are most closely related to the Western North American blackberry species *R. ursinus* [6].

*Rubus* plants are unusual among fruit crops because they typically have perennial crowns and root systems and biennial canes. First-year canes, which are usually vegetative, are called primocanes, while second-year canes that have overwintered are called floricanes. Floral initiation typically begins on primocanes in short days during the autumn, with flowers and fruits developing on floricanes the following spring [7–9]. Raspberry and blackberry cultivars with this customary flowering trait are described as floricane-or biennial-fruiting. Primocane- or annual-fruiting red raspberry cultivars that initiate flowers in the early summer and produce fruit on the tip portion of primocanes or primocane branches during the late summer and autumn (Figure 1) were first developed in the 1950s and 1960s [10], with primocane-fruiting blackberries first commercially released in the early 2000s [11]. Primocane-fruiting cultivars differ from traditional floricane-fruiting types in that they have no short-day requirement for flower induction and low-temperature requirement for flower emergence. Primocane-fruiting raspberries and blackberries have grown in economic importance over the past two decades because they confer several advantages for growers. The primocane crop is typically distinctly later than the floricane crop, which allows for season extension and the possibility for ‘double-cropping’ by producing a floricane crop followed by a primocane crop from the same plant in each year. Furthermore, primocane-fruiting allows for production in an expanded geographical area, including tropical areas where there would be insufficient chilling hours for floricane cultivars, and regions where winter injury to canes is problematic [12].

**Figure 1.**
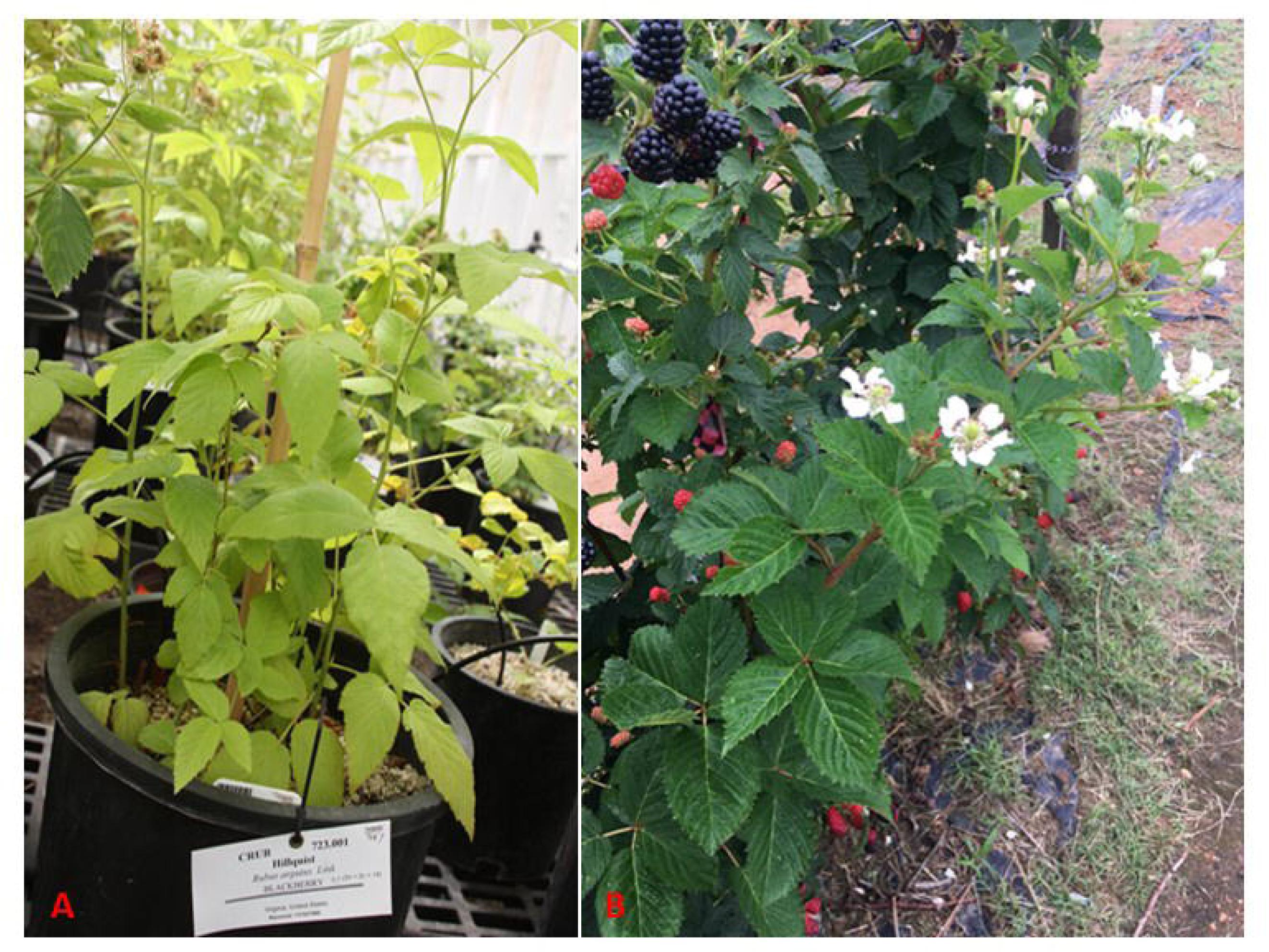
(A) Hillquist blackberry (PI 553951) at the USDA National Clonal Germplasm Repository and (B) a primocane-fruiting blackberry with ripe fruit on second-year canes (floricanes) and flowers on the tip of a first-year cane (primocane).

The only known source of primocane-fruiting in tetraploid blackberry cultivars is a recessive allele from the wild diploid accession ‘Hillquist’ (*R. argutus*; PI 553951; Figure 1). ‘Hillquist’ was initially discovered in Ashland, VA by L.G. Hillquist, who noticed that some of the wild blackberries growing in his backyard had an unusual fruiting habit. The accession was later donated to the New York State Agricultural Experiment Station by Mrs. Hillquist in 1949 [13]. ‘Hillquist’ was first used as a male parent in crosses with the tetraploid, floricane-fruiting blackberry cultivar ‘Brazos’ in 1967, but the first primocane-fruiting cultivars, ‘Prime-Jim’^®^ and ‘Prime-Jan’^®^ were not released until nearly 40 years later [11, 12]. Since then, many public and private blackberry breeding programs have accessed this germplasm, and ‘Hillquist’ is in the pedigree of many important floricane-fruiting and primocane-fruiting cultivars grown around the world.

Despite their economic importance, very few genomic resources are available for blackberries compared to other fruit crops. Pseudo-chromosome level genome assemblies are available for over 20 Rosaceae crops, including apple (*Malus × domestica*) [14–16], peach (*Prunus persica*) [17], and Asian pear (*Pyrus pyrifolia*) [18]. Within the *Rosoideae* subfamily, which is characterized by a base chromosome number of x=7, there are high-quality genome assemblies available for rose (*Rosa chinensis*) [19] and diploid (*Fragaria vesca*) [20, 21] and octoploid (*F. × ananassa*) [22] strawberry. The first *Rubus* genome sequenced was a highly homozygous diploid black raspberry selection, ORUS 4115-3 [23–25]. More recently, chromosome-length assemblies have been published for the red raspberry cultivar ‘Anitra’ [26] and *R. chingii* [27]. To date, however, the only published *Rubus* genome assemblies are for species in subgenus *Ideaobatus,* and there is no genome sequence data for any close relatives of cultivated blackberries in subgenus *Rubus* in public databases.

Here we present a chromosome-length genome assembly and annotation of the diploid *R. argutus* accession ‘Hillquist’. ‘Hillquist’ was chosen for the assembly because it is the original source of the primocane-fruiting used in cultivated tetraploid blackberries and is now represented in the pedigree of public and private blackberry breeding germplasm around the world. The *R. argutus* assembly was produced using PacBio long-read single-molecule real-time (SMRT) sequencing and scaffolded using Hi-C sequence data. The full assembly is 298.2 Mb in length, with 270.0 Mb (90.1%) assigned to seven scaffolds with an average length of 38.6 Mb. Repetitive elements were predicted to make up 52.8% of the genome, with *Gypsy* superfamily lineages accounting for the largest fractions of LTR-REs. The computational annotation was performed with support of RNA-seq and Iso-seq data generated from root tips and actively growing leaves and stems of primocane and floricanes. A total of 38,503 protein-coding genes were predicted from the genome, 72.2% of which were functionally annotated. The practical value of the *R. argutus* genome assembly and annotation was demonstrated by comparing the genome sequences of related Rosoideae species [19,21,25–27], anchoring the scaffolds to a novel modified genotyping-by-sequencing-based (GBSpoly) linkage map of tetraploid blackberry, and identifying possible candidate genes for primocane-fruiting within a previously mapped locus.

### Plant material

The *R. argutus* germplasm accession ‘Hillquist’ (PI 553951), sourced from the USDA National Clonal Germplasm Repository (NCGR) was used for genome sequencing and assembly. Leaf material was harvested from a single plant of this cultivar grown in the greenhouse at the USDA-NCGR, in Corvallis, Oregon for flow cytometry, DNA extraction, and PacBio, 10x Chromium, and Hi-C sequencing. ‘Hillquist’ plants propagated by NCGR staff were sent to North Carolina State University (NCSU) and grown in a greenhouse. Tissue from root tips and actively growing leaves and stems from primocanes and floricanes for RNA sequencing and IsoSeq was obtained from plants grown at NCSU.

### Genome size estimation

Nuclear flow cytometry with DAPI staining was used to measure DNA content and estimate the genome size of *R. argutus* ‘Hillquist’. Flow cytometry was performed using young, unexpanded ‘Hillquist’ leaves in biological triplicate with *Vinca major* as an internal standard. The nuclear flow cytometry generated estimate of the *R. argutus* genome size was 337.4 Mb (1C = 0.345 pg). This estimate falls within the reported range of other diploid species in subgenus *Rubus* (*R. hispidus*, *R. canadensis*, *R. trivialis*, *R. canescens*, and *R. sanctus*), which was between 1C = 0.295 – 0.375 pg [28, 29].

### DNA extraction, library preparation, and sequencing

#### PacBio

High molecular weight DNA was extracted from young, unexpanded leaves of *R. argutus ‘*Hillquist’ using a modified CTAB method [30]. DNA quality was evaluated with Pulsed Field Gel Electrophoresis (BioRad, Hercules, California), and quantification was performed with a Qubit fluorometer (ThermoFisher Sci., Waltham, Massachusetts). Genomic DNA was sheared to achieve fragments in the 15–40 kb size range using a 26 gauge blunt end needle (ThermoFisher UK Ltd HCA-413-030Y GC Syringe Replacement Parts 26 g, 51 mm) and 1 mL luer-loc syringe. The sheared DNA was then cleaned using 1X AMPure PB Beads before library preparation. Fragments were enzymatically repaired and used to construct a long read (20 kb) PacBio Sequel genomic library with a SMRTbell™ Template Prep Kit 1.0-SPv3 according to the manufacturer’s recommendations (Pacific Biosciences Inc., Menlo Park, CA, USA). The resulting SMRTbell templates were size selected using BluePippin electrophoresis (Sage Science Inc., Beverly, MA, USA) and template DNA ranging in size between 15 and 50 kb was sequenced in eight PacBio Sequel Single-Molecule Real-Time (SMRT) cells on a PacBio Sequel instrument at the NCSU Genomic Sciences Laboratory. A combined total of 3.8 million PacBio post-filtered reads with an average length of 6,824 bp were generated from the eight SMRT cells, resulting in a total of 25.9 Gb of sequence (∼77X Genome Coverage) (Supplemental Table S1). *Hi-C and 10x Genomics*

Five grams of young leaf tissue for Hi-C and 10x Genomics library preparation was collected from a ‘Hillquist’ plant subjected to 48 hours of darkness. An *in situ* Hi-C library was prepared following [31] and sequenced to produce 559,559,351 paired-end reads. 10x Genomics linked read libraries were made at the Wellcome Sanger Institute High-Throughput DNA Sequencing Centre by the Sanger Institute R&D and pipeline teams using the Chromium^TM^ Genome Reagent Kit (v2 Chemistry) following the manufacturer’s recommended protocol. These libraries were then sequenced on Illumina NovaSeq 6000 platforms at the Wellcome Sanger Institute High-Throughput DNA Sequencing Centre.

### Genome sequence assembly

A contig-scale assembly was generated with PacBio sequence data using the FALCON and FALCON-Unzip software applications [32]. Error correction on the phased assembly was performed using the Arrow consensus model in the PacBio GenomicConsensus package following default parameters. The FALCON-Unzip assembly comprised 374 Mb of sequence in 1756 contigs with an N50 of 486 kb and a maximum contig length of 5.9 Mb. The K-mer distribution of unassembled, corrected PacBio reads for ‘Hillquist’ showed a bimodal distribution, indicating high heterozygosity. Therefore, the Purge Haplotigs pipeline was used to curate the heterozygous diploid genome assembly and resolve under-collapsed heterozygosity by identifying syntenic pairs of contigs and moving one to a haplotig pool [33]. The optimized Purge Haplotigs assembly consisted of 297 Mb assigned to 811 primary contigs with a contig N50 of 650 Kb and a maximum contig length of 5.9 Mb.

Hi-C data were aligned to the Purge Haplotigs draft assembly using Juicer v1.6.2 [34]. A candidate assembly and contact maps visualizing the alignments with respect to the draft and the new reference were built using the 3D *de novo* assembly (3D-DNA) pipeline [35], and the genome was reviewed and polished using Juicebox Assembly Tools [36]. The new 298 Mb assembly was composed of 350 scaffolds with an N50 of 38.6 Mb and a maximum scaffold length of 45.5 Mb (Table 1; Figure 2). Among these Hi-C scaffolds, seven chromosome-length scaffolds with a total length of 270 Mb (90% of the 298 bp genome) corresponded directly to the seven *R. occidentalis* and *F. vesca* chromosomes (Supplemental Table S2). Chromosome nomenclature and orientation were assigned following *Fragaria* conventions [20].

**Figure 2.**
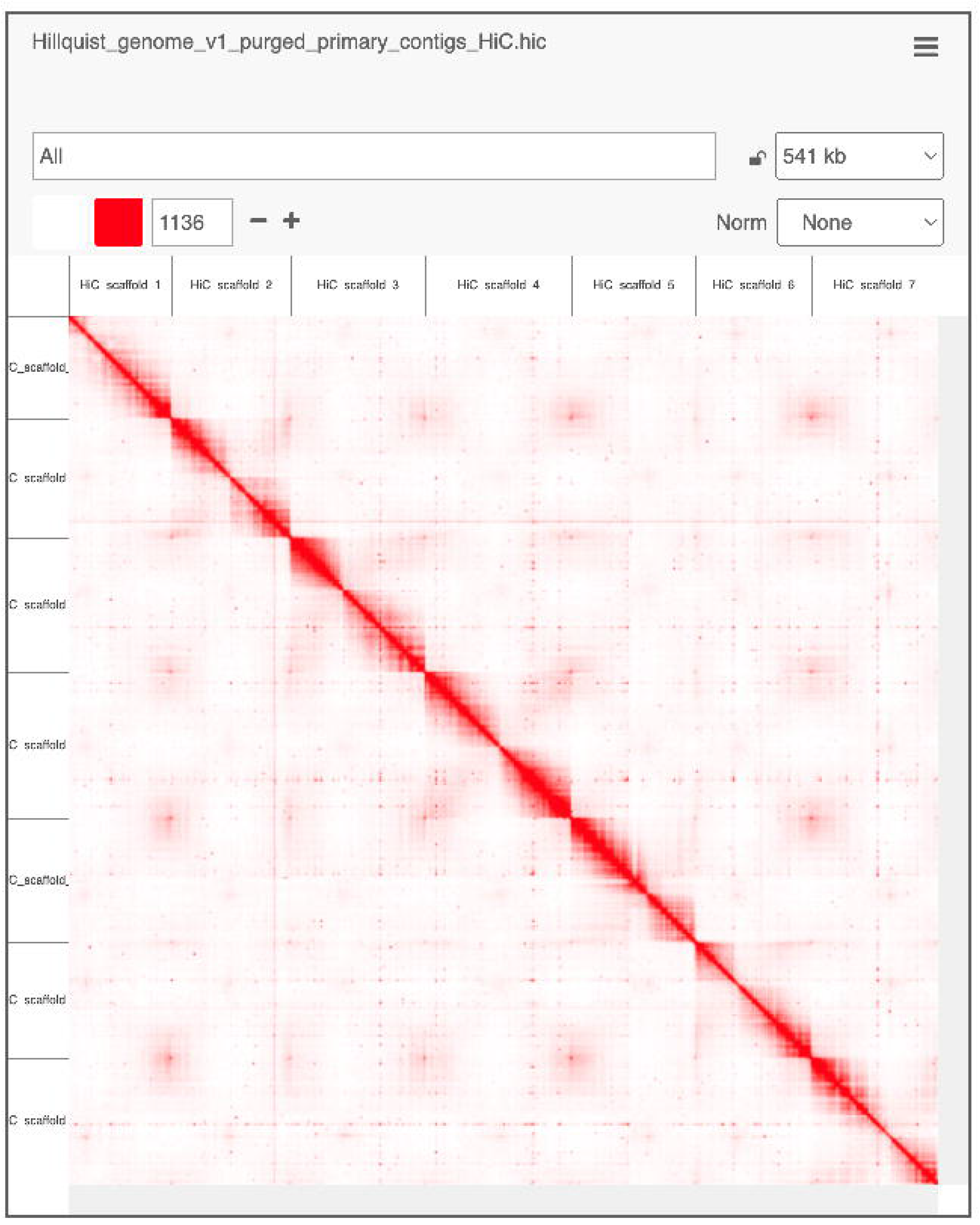
Hi-C interaction matrix for the ‘Hillquist’ blackberry (*R. argutus*) assembly. An interactive version of this map is available at https://tinyurl.com/y7vvu9mc.

**Table 1.**
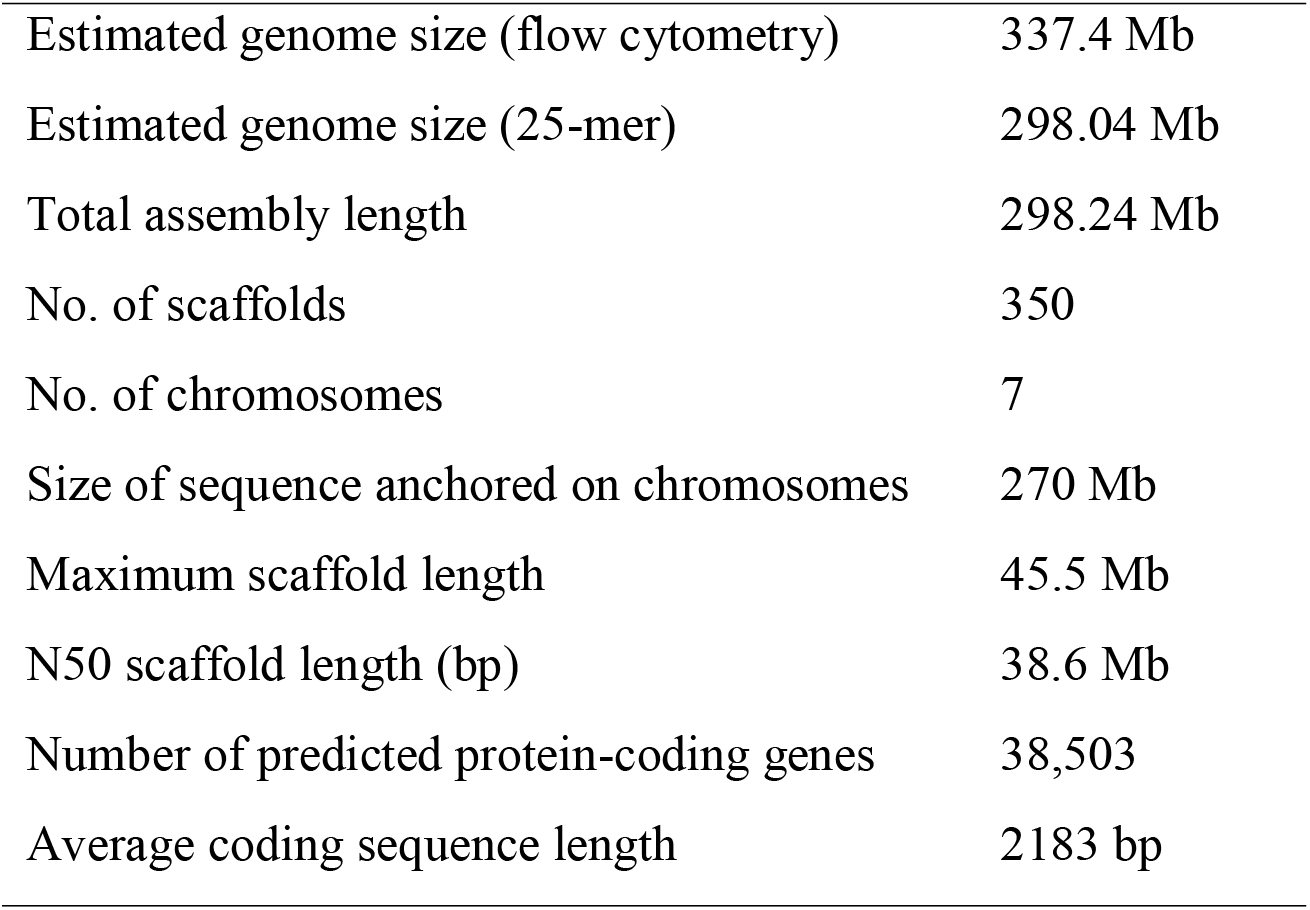
Summary statistics for the assembled *R. argutus* ‘Hillquist’ genome.

Heterozygosity and genome size were estimated by analysis of the *k-mer* count histogram generated with 10x Chromium Illumina reads using the online version of GenomeScope (GenomeScope, RRID:SCR_017014; [37]). The size and heterozygosity of the genome were estimated as 298.06 Mb and 1.04% (Supplemental Figure S1). The *k-mer* based genome size estimate was within 172.4 kb of the Hi-C assembly length, which suggests that the genome was nearly complete. However, the flow cytometry estimate of *R. argutus* genome size was 337 Mb, indicating that 88.4% of the genome was incorporated in the assembly.

### Synteny with Rosoideae genomes

Synteny of the ‘Hillquist’ genome to the other publicly available Rosoideae genome sequences (*R. idaeus* ‘Anitra’ [26], *R. chingii* [27], *R. occidentalis* [25], *F. vesca* ‘Hawaii 4’ [21], and *Rosa chinensis* ‘Old Blush’ [19]) was determined with MUMmer4 [38] using default parameters. Data for the genomes were downloaded from the data repository on the Genome Database for Rosaceae (https://www.rosaceae.org; [39]), and the associations revealed were plotted using R following Davik et al. [26]. The ‘Hillquist’ assembly showed a high degree of collinearity to the other Rosoideae genomes (Figure 3, Supplementary Figure S2). Collinearity to the genomes of *R. idaeus* ‘Anitra’ and *R. chingii* was particularly high, with no large-scale rearrangements, translocations, or inversions observed across any of the seven chromosomes when compared to these two species (Figure 3c, Supplementary Figure S2). As with the previously published comparison of the *R. idaeus* ‘Anitra’ and *R. occidentalis* genome assemblies [26], several areas of non-collinearity were observed between the ‘Hillquist’ and *R. occidentalis* genomes. The most notable areas of non-collinearity between the genomes were the large degree of rearrangement on one half of chromosomes 1 and 4 and the large inversions observed on chromosome 6 (Figure 3d). Other authors [26, 40] have suggested that these differences could be the result of errors in the assembly of the *R. occidentalis* genome, and the data presented here support that hypothesis. The pattern of rearrangements observed between the ‘Hillquist’ genome and *F. vesca* ‘Hawaii 4’ and *R. chinensis* ‘Old Blush’ genomes were similar to those previously reported for the ‘Anitra’ genome [26]. Two large inversions on chromosomes 5 and chromosome 7 and with several smaller inversions on chromosomes 3 and 4 were observed between the ‘Hillquist’ genome and that of *F. vesca* ‘Hawaii 4’ (Figure 3a). Two significant translocations were observed between chromosome 1 and chromosome 6 of the ‘Hillquist’ and *R. chinensis* ‘Old Blush’ genomes, along with small inversions on chromosomes 2 and 7 (Figure 3b). These rearrangements reflect the evolutionary timescales since the *Rubus*, *Fragaria,* and *Rosa* ancestral genomes diverged from a common ancestor [41].

**Figure 3.**
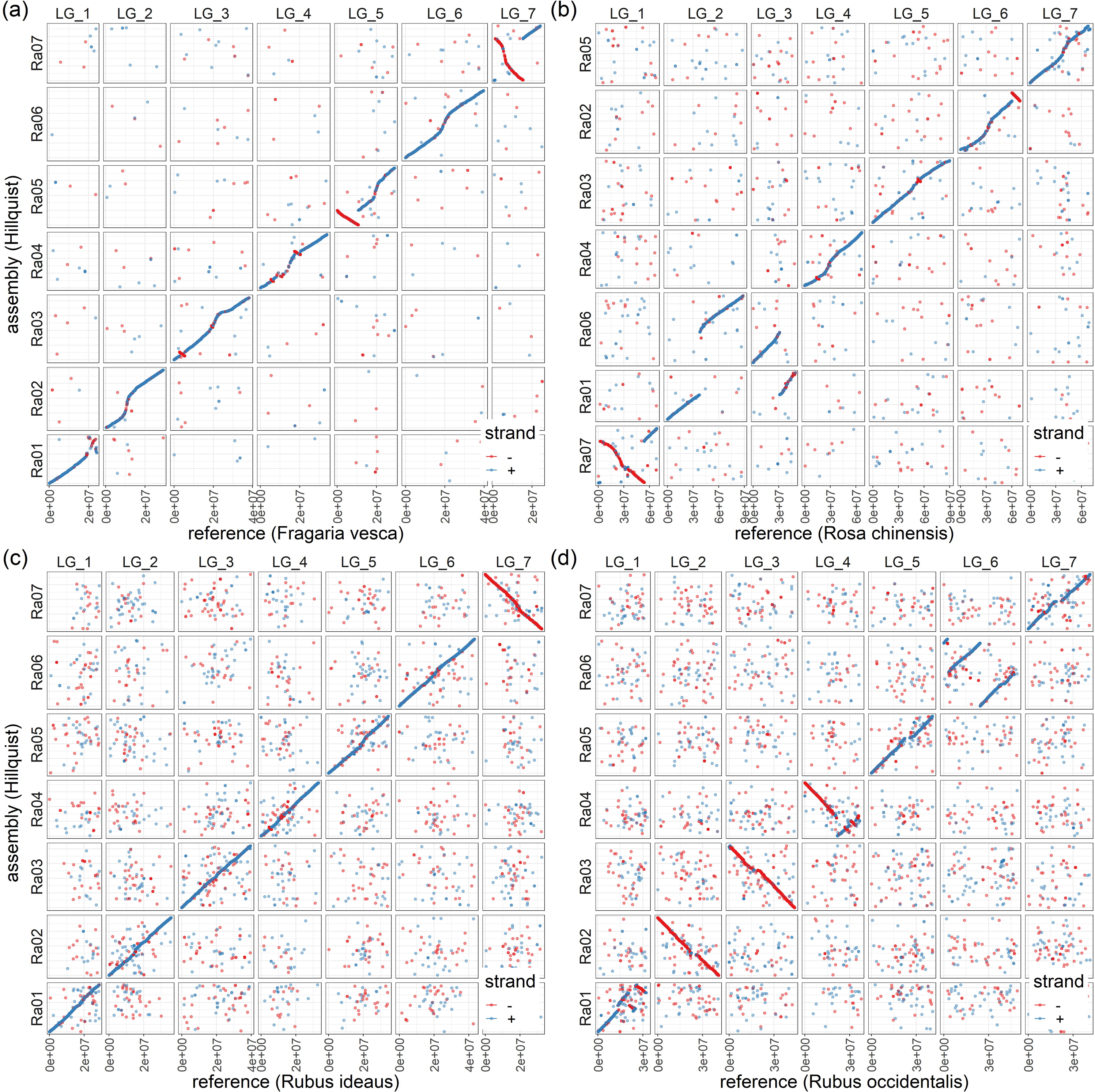
Whole-genome alignment plots between the ‘Hillquist’ blackberry (*R. argutus*) genome assembly and the chromosome-length assemblies of (a) woodland strawberry (*Fragaria vesca* V. 4), (b) rose (*Rosa chinensis*), (c) red raspberry (*R. idaeus*), and (d) black raspberry (*R. occidentalis* v.3).

### Linkage Map of Autotetraploid Blackberry

The best available blackberry linkage map was constructed using 119 simple sequence repeat markers developed from red raspberry and a blackberry expressed sequence tag library [42]. Due to the paucity of markers, this SSR-based map contained large genetic regions with no marker coverage. The utility of the ‘Hillquist’ genome sequence for use in fresh-market blackberry breeding was therefore assessed by anchoring the pseudo-chromosomes to a novel linkage map of the autotetraploid breeding selection, A-2551TN, from the University of Arkansas System Division of Agriculture Fruit Breeding Program. The linkage map consisted of 2,935 sequence-characterized markers that were identified using a modified genotyping-by-sequencing protocol (GBSpoly) that is robust for highly heterozygous and polyploid genomes. Full methods and results for map construction are provided in Supplemental File 1. Briefly, a mapping population consisting of 119 F_1_ progeny from the cross A-2551TN x APF-259TN (Supplemental Figure S3) were used to generate a maternal haplotype map. Multiplexed NGS-based reduced representation sequencing libraries for parents and progeny were prepared following the GBSpoly protocol optimized for heterozygous and polyploid genomes [43, 44] and sequenced on the HiSeq 2500 (Illumina, San Diego, CA) and the SP flow cell of the NovaSeq™ 6000 (Illumina, San Diego, CA) system at the Genomic Sciences Laboratory at NCSU to generate 615.4 million sequencing reads after demulitplexing and quality filtering. Raw Fastq files were processed and filtered with the ngsComposer [45] pipeline (https://github.com/bodeolukolu/ngsComposer) and were aligned to the black raspberry [25] and ‘Hillquist’ genomes using BWA-MEM [46]. The GBSapp pipeline (https://github.com/bodeolukolu/GBSapp), which integrates original and third-party tools (bwa, samtools, picard, bcftools, GATK, java, R-ggplot2, and R-AGHmatrix), was used for variant calling and filtering. In total, 85.9% of reads were mapped to unique positions on the ‘Hillquist’ genome, while only 67.3% of reads mapped to unique positions on the black raspberry genome (Supplemental Table S3). A total of 1,811,617 and 2,022,664 polymorphic markers were identified when these reads were aligned to the black raspberry and ‘Hillquist’ genomes, respectively.

Only single dose markers segregating in A-2551TN were used to construct the haplotype-resolved maternal linkage map. After quality filtering in the GBSapp pipeline [43] and extracting only single dose markers, 3,796 markers remained that had less than 5% missing data, were heterozygous in A-2551TN (0/0/0/1 x 0/0/0/0), and segregated in a 1:1 ratio in the progeny. A maternal haplotype map composed of 2,935 markers assigned to 30 linkage groups, with between 5 and 249 markers per linkage group, was then created using JoinMap 4.1 [47]. The total map length was 2,411.81 cM, with linkage groups ranging from 18.61 cM to 146.65 cM in length and an average of one marker every 0.82 cM (Supplemental Tables S4, S5; Supplemental Figure S4). The physical positions of the mapped markers on the ‘Hillquist’ pseudo-chromosomes were used to identify four homologous linkage groups corresponding to five chromosomes (1, 2, 3, 4, and 6), and five homologous linkage groups corresponding to the remaining two chromosomes (5 and 7). The A-2551TN maternal haplotype map was strongly collinear with the ‘Hillquist’ genome, with no major translocations or inversions (Figure 4). While many of the linkage groups in the A-2551TN maternal haplotype map contained markers that aligned to physical positions across the length of each of the chromosomes, ten linkage groups had markers aligned to physical positions spanning less than 10 Mbp in the ‘Hillquist’ genome. Based on the physical positions of these markers on short linkage groups, it is likely that linkage groups 7b/7d and 5c/5e belong to the same haplotype of A-2551TN. Gaps in the linkage map can likely be attributed to the high inbreeding coefficients of A-2551TN (F= 0.100) and its progeny from the A-2551TN x APF-259TN cross (F = 0.099). The high percentage of reads mapped to unique positions on the ‘Hillquist’ genome and the collinearity between the physical map of ‘Hillquist’ and the A-2551TN maternal haplotype map validate the order and orientation of the Hi-C-based chromosome-length assembly of ‘Hillquist’ and demonstrate its utility for genomic breeding research in polyploid fresh-market blackberries.

**Figure 4.**
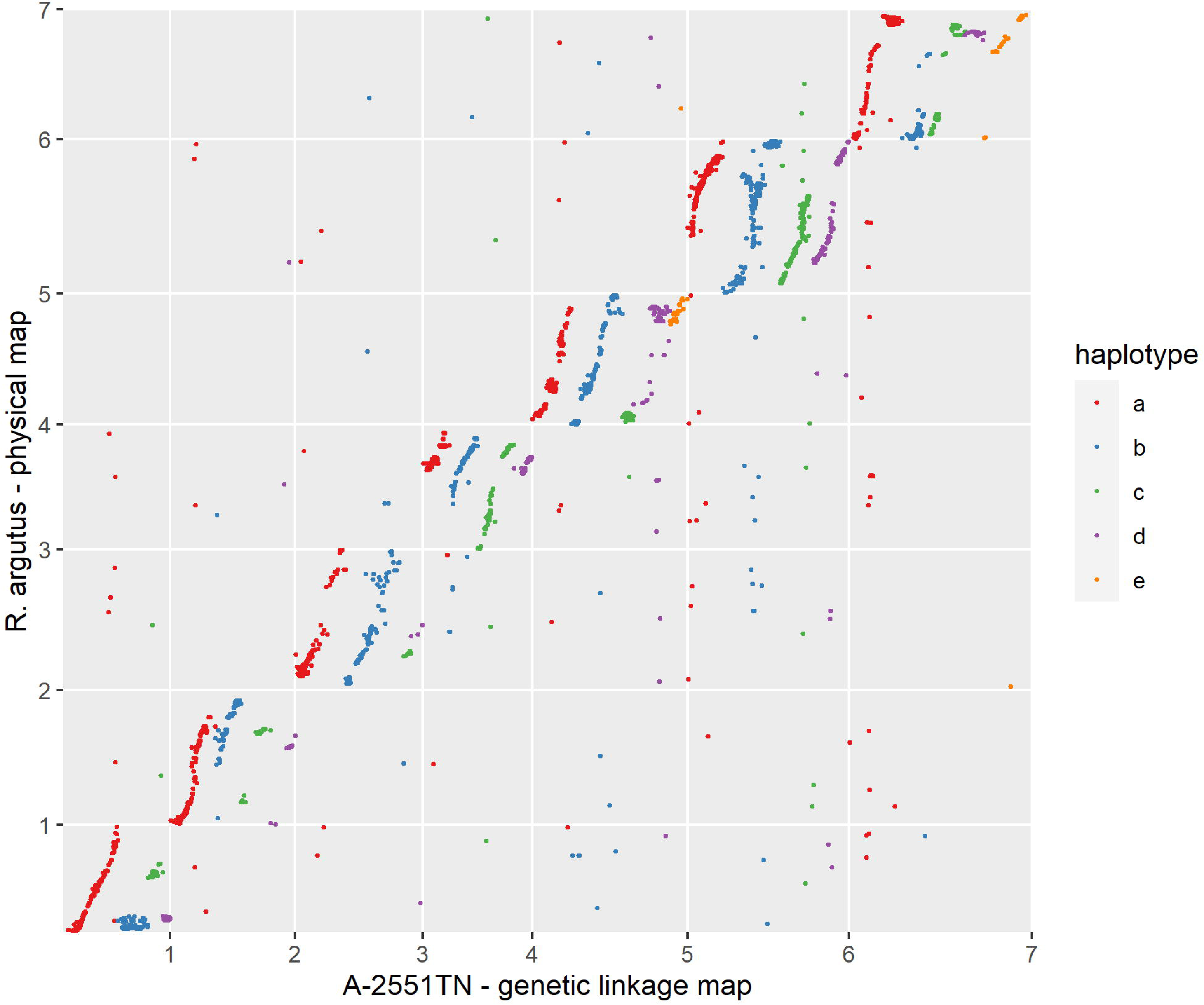
Comparison of the tetraploid A-2551TN maternal haplotype map with the ‘Hillquist’ blackberry (*R. argutus*) physical map. As expected, four homologous linkage groups (haplotypes a-d) were identified for chromosomes Ra01, Ra02, Ra03, Ra04, and Ra06. Five homologous linkage groups (haplotypes a-e) corresponded to chromosomes Ra05 and Ra07. Based on the physical positions of the markers on chromosomes Ra05 and Ra07, it is likely that linkage groups 5c and 5e and 7b and 7d actually belong to the same haplotype of A-2551TN.

### Analysis of Repetitive Content

The repetitive component of the ‘Hillquist’ genome was analyzed using both structural-and clustering-based characterization analyses. Structural-based results were compared with those of the other four Rosaceae species (*F. vesca*, *Potentilla micrantha*, *P. persica* and *M. domestica*). The data of the other four Rosaceae species were retrieved from the NCBI database (NCBI, Washington, USA, https://www.ncbi.nlm.nih.gov/) and the GigaScience GigaDB repository (Supplemental Table S6). The quality of the ‘Hillquist’ paired-end Illumina reads was inspected using FastQC v0.11.5 [48], and Illumina adapters and low-quality regions were removed using Trimmomatic v0.39 [49] with the following parameters: ILLUMINACLIP:2:30:10; LEADING:3; TRAILING:3; SLIDINGWINDOW:4:15; CROP:90; MINLEN:90. Duplicated reads were removed using the prinseq-lite.pl script v0.20.4 with -derep 1 [50]. Organellar sequences were removed from the datasets by mapping against an ad hoc prepared set of chloroplast genomes of *F. vesca* (NCBI JF345175.1), *M. domestica* (NCBI MK434916.1), *P. micrantha* (NCBI HG931056.1), *P. persica* (NCBI HQ336405.1) and *Rubus leucanthus* (NCBI MK105853.1) and mitochondrial genomes of *M. domestica* (NCBI NC_018554.1) and *Prunus avium* (NCBI MK816392.2) using CLC-BIO Genomic Workbench v9.0.4 (CLC-BIO, Aarhus, Denmark) with the following parameters: mismatch cost 1; insertion cost 1; deletion cost 1; length fraction 0.9; similarity fraction 0.9. All matching sequences were considered putatively belonging to organellar genomes and subsequently removed.

#### Clustering-based characterization of repeats

A clustering characterization of the repetitive component of the ‘Hillquist’ genome was performed using RepeatExplorer2 [51] with default parameters with a random set of 1,000,000 paired-end sequences. To reduce the number of unknown retrotransposon clusters, BLASTN and tBLASTX [52] analyses were performed using BLAST v2.6.0 with default parameters against the libraries of the characterized Rosaceae full-length long terminal repeat retrotransposons (LTR-REs). Of the 1 million ‘Hillquist’ paired-end Illumina reads randomly selected for *de novo* clustering, 555,442 reads were processed by RepeatExplorer2. Of these processed reads, 262,064 (47.2% of the genome) were considered singlets and did not fall into the category of repeated sequences according to the thresholds imposed by the program. The remaining 293,379 reads (52.8% of the genome) were characterized as repeats and grouped in 51,851 clusters, each of which represented a single repeat sub-lineage. One hundred and seventy-three clusters were classified as top clusters with a genome proportion greater than 0.01%, representing the most abundant repeat families. *Copia* and *Gypsy* superfamilies accounted for the largest fractions of the genome (10.51% and 23.44%, respectively; Table 2). In particular, *Athila*-related clusters were the most abundant. No DNA transposons, non-LTR elements, or satellite DNA were among the top clusters. The absence of DNA transposon and satellite DNA in top clusters indicates that these repeat types are scarce in the *Rubus* genome. Illumina reads related to these repeats were assembled in clusters accounting for less than 0.01% of the genome. Finally, 18.14% of the repetitive component remained unclassified.

**Table 2.**
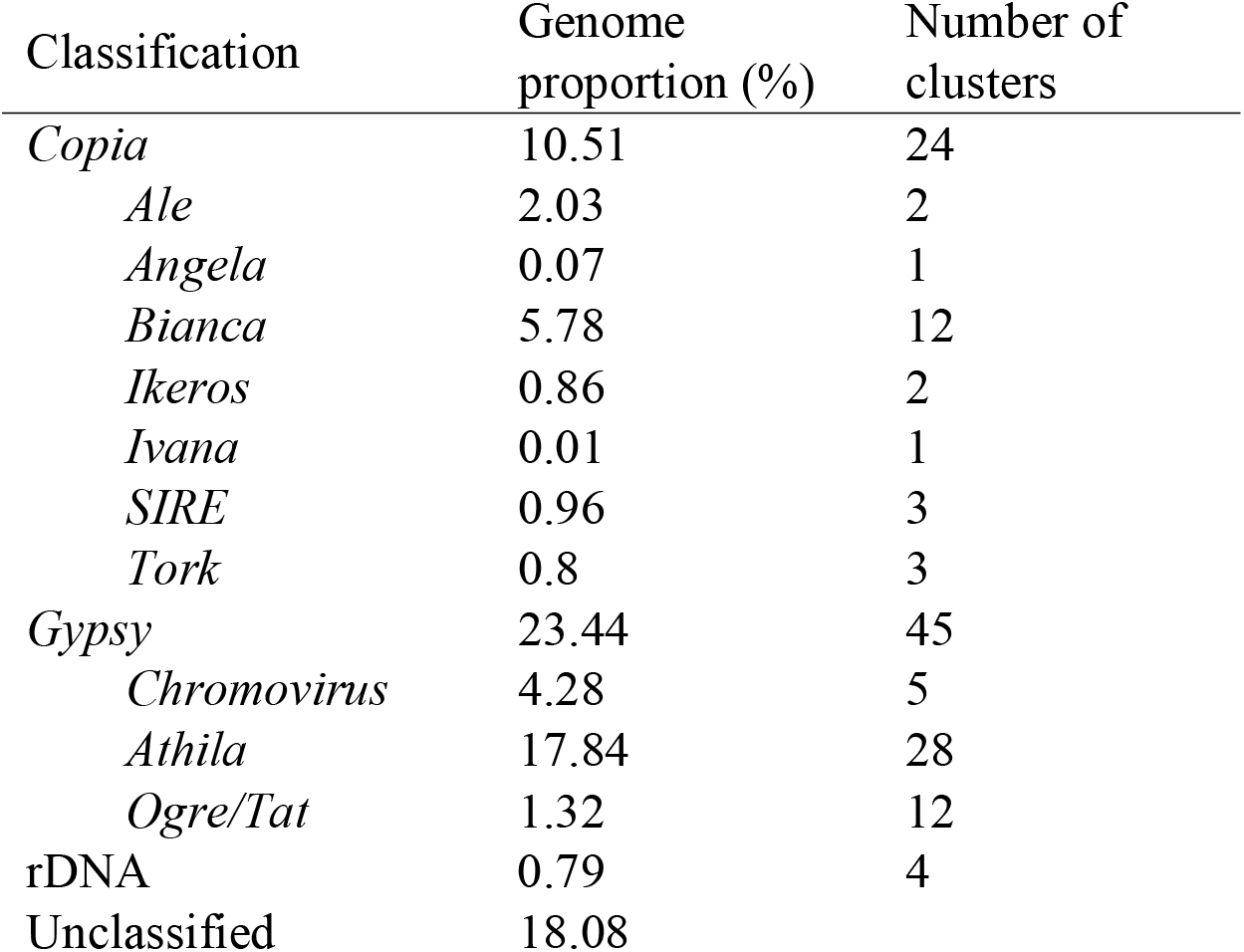
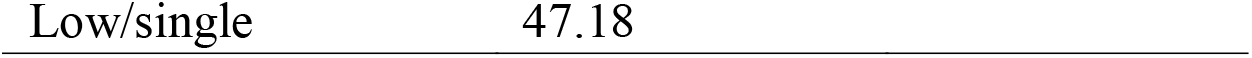
Classification of clusters produced by RepeatExplorer2 and proportion of repeat types in the genome of ‘Hillquist’ (*Rubus argutus*).

| Classification | Genome proportion (%) | Number of clusters |
| --- | --- | --- |
| *Copia* | 10.51 | 24 |
| *Ale* | 2.03 | 2 |
| *Angela* | 0.07 | 1 |
| *Bianca* | 5.78 | 12 |
| *Ikeros* | 0.86 | 2 |
| *Ivana* | 0.01 | 1 |
| *SIRE* | 0.96 | 3 |
| *Tork* | 0.8 | 3 |
| *Gypsy* | 23.44 | 45 |
| *Chromovirus* | 4.28 | 5 |
| *Athila* | 17.84 | 28 |
| *Ogre/Tat* | 1.32 | 12 |
| rDNA | 0.79 | 4 |
| Unclassified | 18.08 |  |

#### Full-length LTR-retrotransposon discovery and characterization analysis

The genome assemblies of ‘Hillquist’*, F. vesca*, *P. micrantha*, *P. persica*, and *M. domestica* were scanned for a structural identification of Class I full-length LTR-REs using LTRharvest v1.5.10 [53] with the following parameters: -minlenltr 100; -maxlenltr 6000; - mindistltr 1500; -maxdistltr 25000; -mintsd 5; -maxtsd 5; -similar 85; -vic 10; -motif tgca. The libraries of full-length LTR-REs were submitted to domain-based annotation by using DANTE v1.0.0, available on the RepeatExplorer Galaxy-based website (https://repeatexplorer-elixir.cerit-sc.cz/galaxy/). The annotation process was performed with default parameters using the REXdb of transposable element protein domains [54] and a BLOSUM80 scoring matrix. The protein matches were filtered by significance using the parameters provided by the platform, and nested elements were manually removed. To reduce the number of uncharacterized full-length LTR-REs, we performed BLASTN and tBLASTX between uncharacterized elements and characterized elements in conjunction with the annotated contigs produced by the comparative clustering analysis.

A total of 636 full-length LTR-REs were identified in the ‘Hillquist’ genome assembly, with 217 and 409 LT-REs belonging to the *Gypsy* and *Copia* superfamilies, respectively (Supplemental Table S7). The number of full-length LTR-REs isolated from the other four genome assemblies of Rosaceae species varied from a minimum of 204 in *F. vesca* to a maximum of 2,662 in *M. domestica* (Supplemental Table S7). *Copia* elements were more abundant than *Gypsy* LTR-REs in ‘Hillquist’ (1.9:1), *F. vesca* (2.7:1) and *P. persica* (4.5:1), while LTR-REs in *Copia* and *Gypsy* superfamilies were equally represented in *P. micrantha* (1:1), and *Gypsy* elements were slightly more abundant in *M. domestica* (0.7:1). The lineage level annotation of most elements revealed considerable quantitative and qualitative variability among the five species, with several lineages that were not detected in some species. However, it is possible that very ancient and rearranged elements may not have been identifiable through structural features due to the stringency of the parameters used in the identification process.

### Gene prediction and annotation

#### RNA extraction, library preparation, and sequencing

Total RNA was extracted from five tissue types (root tips, as well as actively growing leaves and stems from both primocane and floricane canes of the same plant) for sequencing with RNA-Seq and Iso-Seq technologies using the Spectrum Plant Total RNA Kit (Millipore Sigma, Burlington, MA) following the manufacturer’s protocol. Cane types were distinguished by the presence of trifoliate leaves on floricanes and pentifoliate leaves on primocanes. The purity and concentration of the extracted RNA was determined using a 2100 Bioanalyzer (Agilent Technologies, Santa Clara, CA), and the integrity of the samples was determined using a Qubit 4.0 fluorimeter (Thermo Fisher Scientific, Waltham, MA). Samples with an RNA integrity number (RIN) value above 7.0 were submitted for subsequent sequencing. Two duplicate RNA-Seq libraries were produced for each tissue type and sequenced with an Illumina HiSeq X instrument at Scientific Operations core at the Wellcome Sanger Institute. A total of 135,518,570 paired reads were generated from the 10 RNA-Seq libraries, with 9,457,856 to 20,119,374 paired reads per library.

Total RNA from the same five tissue samples were pooled and used for Iso-Seq library preparation. Standard PacBio Iso-Seq SMRTbell libraries were prepared by Genewiz (South Plainfield, NJ). In total, one SMRT cell was sequenced with Sequel II to generate a total of 5,959,439 polymerase reads were generated with a mean length of 39,878 bp per read, an average insert length of 7,387 bp, and a mean subread length of 1,614. Full-length transcripts were identified using the Iso-Seq 3 application in SMRTLink 5.0. First, multiple reads of the same SMRTbell sequence or the subreads from the same polymerase read were combined to produce one high-quality circular consensus sequence (CCS). A total of 2,830,415 CCS reads with a mean length of 1,526 bp were generated in this process. Next, the CCS reads were classified as full-length based on the presence of both cDNA primers and polyA tails in the reads. Full-length reads were further classified as chimeric or non-chimeric reads based on whether or not primers were found in the middle of the sequences. Finally, unpolished consensus isoforms were extracted using the iterative clustering and error correction algorithm and polished to obtain high-quality and low-quality isoforms. A total of 185,699 and 290 polished high-quality and low-quality isoforms were generated in this process.

#### Structural Gene Annotation

A repeat library of transposable element families was generated using RepeatModeler2 [55]. Repeat sequences, interspersed repeats, and low complexity DNA sequences were identified and soft-masked using RepeatMasker [56]. Repeat masking was further refined using Iso-Seq transcript sequences. The representative Iso-Seq open reading frames (ORFs) supported by protein or RNA-Seq evidence were used to reduce the amount of repeat-masked coding sequence by unmasking the masked regions overlapping the Iso-Seq defined ORFs, and as such, 135 Mbp (45.4%) of the ‘Hillquist’ genome was repeat masked. The repeat length distribution is shown in Supplementary Figure S5. Masking refinement based on aligned Iso-Seq transcripts unmasked 1.9 Mbp of the sequence at 7,257 distinct Iso-Seq loci. RNA-Seq mapping originated intron hints were obtained by aligning paired RNA-Seq reads to the ‘Hillquist’ genome using STAR [57] with filters for the intron coverage value ≥ 3. Additionally, consensus high-quality Iso-Seq isoforms were aligned to the genome by GMAP [58] with filters for ≥ 95% identity and ≥90% coverage. The longest open reading frame (lORF) was identified in each aligned transcript. In loci with overlapping isoforms, a single representative transcript with the longest lORF was selected thus making a set of non-overlapping Iso-Seq isoforms. Transcripts with lORFs shorter than 300 nt or with introns longer than 10k were filtered out from this set. Protein hints to splice sites and translation initiation and termination sites were generated by ProtHint [59] using proteins from the Plantae section of the OrthoDB v10 protein database [60].

Genes were annotated using a protocol similar to BRAKER2 [61], with additional integration of RNA-Seq and Iso-Seq data (Supplemental Figure S6). GeneMark-ET [62] with RNA-Seq intron hints was used to create a set of predicted genes. In this analysis, introns mapped with coverage ≥100 were used for initial parameter estimation. Genes predicted by GeneMark-ET were subsequently used as seed regions in ProtHint to generate protein hints. Next, protein and RNA-Seq hints were used together to predict genes with GeneMark-EP+ [59]. By default, GeneMark-EP+ directly uses protein hints generated by ProtHint. This hint set was extended by adding RNA-Seq intron hints. Introns found in the intersection of RNA-Seq and protein hints were added to GeneMark-EP+’s high-confidence hint set. Genes predicted by GeneMark-EP+ and ORFs from the set of non-overlapping Iso-Seq isoforms were combined to create the new seed regions. In case of an overlap between the Iso-Seq and GeneMark-EP+ defined seeds, the Iso-Seq seed was selected if its ORF was > 50 nt longer than the GeneMark-EP+ seed. GeneMark-EP+ was then run on the genome with updated repeat-masking and protein hints delivered by the second iteration of ProtHint. Again, RNA-Seq hints were added to the hints set in the same way as described for the first GeneMark-EP+ run. GeneMark-EP+ predictions fully supported by mapped Iso-Seq transcripts or protein hints were selected for the training of AUGUSTUS [63]. AUGUSTUS was run on the ‘Hillquist’ genome sequence with refined repeat-masking and ProtHint proteins hints in agreement with the BRAKER2 protocol [61] to generate the final gene predictions.

The predicted genes were categorized according to their support by external evidence. Multi-exon transcripts were *fully supported* by Iso-Seq if all introns had support by at least a single Iso-Seq transcript. The supporting Iso-Seq transcript could not contain any additional introns, except in its 5’ and 3’ UTRs. Multi-exon transcripts were fully supported by proteins or RNA-Seq if all their introns were supported by protein or RNA-Seq hints. Single exon transcripts were fully supported by Iso-Seq if a matching longest open reading frame was found in one of the Iso-Seq transcripts. Single-exon transcripts were fully supported by proteins if the start and stop codons were supported by protein hints. Transcripts *supported* by any evidence were required to have a part of their gene structure supported by an Iso-Seq, RNA-Seq, or a protein hint.

The final set of predicted genes contained 38,503 coding genes and, with counting alternative isoforms, 40,397 coding transcripts. A total of 13,364 of these transcripts were fully supported by Iso-Seq transcripts, while RNA-Seq data fully supported 13,469 transcripts, and 17,848 transcripts had full protein support (Figure 5a-b); 31,326 transcripts were partially supported by some evidence, and the remaining 9,407 transcripts were pure *ab initio* predictions. Transcripts in the unsupported group were rather short (average protein length 166 AA), with a large fraction (5,129; 55%) lacking any introns. Overall, 19,937 genes had full support from at least one of the external evidence types. The average length of proteins encoded by transcripts with at least one type of external evidence support was 400 AA; this set included 6,125 intronless transcripts. The 38,503 coding genes had an average length of 2,183 bp, containing an average of 3.4 introns per gene and median intron and exon lengths of 152 bp and 132 bp, respectively. Of these coding genes, 36,836 had no alternative isoforms, 1466 had two isoforms, and 201 had three or more isoforms (Supplemental Figure S7). The number of predicted genes in *R. argutus* was comparable to other *Rubus* genomes including *R. idaeus* (39,448 [26]), *R. chingii* (33,130 [27]), and *R. occidentalis* (34,545 [25]).

**Figure 5.**
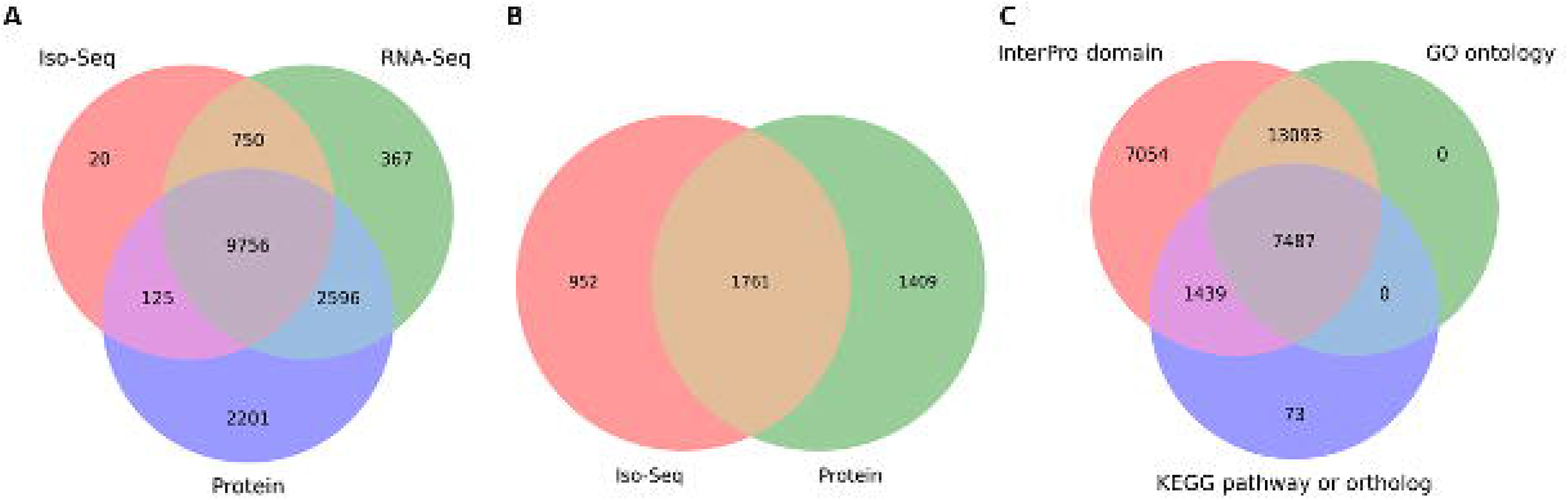
Predicted (A) multi-exon transcripts and (B) single-exon transcripts fully supported by external evidence and (C) predicted transcripts with functional annotation matches.

#### BUSCO analysis

The Benchmarking Universal Single-Copy Orthologs (BUSCO) [64] toolkit was used to assess how many predicted *R. argutus* genes were coding for Universal Single-Copy Orthologs. A total of 2,326 BUSCO gene families defined for the Eudicots odb10 lineage were retrieved. In the predicted set of genes, 2,134 (91.7%) complete *R. argutus* genes orthologous to the BUSCO families were identified, along with 74 (3.2%) genes with partial match. A small fraction of the BUSCO families (5.1%) were not identified among the predicted *R. argutus* genes (Supplemental Figure S8). These results suggest that the ‘Hillquist’ assembly and the gene complement are 94.9% complete.

#### Functional gene annotation

Putative gene function was determined through interrogation of the Swiss-Prot, Araport11, NCBI nr, Refseq and TrEmbl protein databases with BLAST+ blastp-fast algorithm [65] using the predicted protein-coding sequences of the 38,503 genes identified in the structural annotation as queries with an expectation value cutoff of 1e-6. BLAST+ analyses were executed using the Galaxy platform [66] with locally installed databases except for Araport11, which was downloaded from The Arabidopsis Information Resource (TAIR, https://www.arabidopsis.org/). InterProScan v5 [67] was used to assign InterPro domains, and Gene Ontology (GO) terms to the predicted proteins. KEGG ortholog and KEGG pathway mapping were performed with BlastKOALA v2.2 [68] and eggNOG-mapper v2 [69], respectively. Of the 40,397 transcripts predicted in the ‘Hillquist’ genome, a total of 15,333 (37.96%), 22,713 (56.22%), 15,639 (38.71%), 23,370 (57.85%), and 15,986 (39.57%) returned at least one hit after the blastp analysis with nr, Araport11, RefSeq, SwissProt and TrEMBL databases as subjects, respectively (Supplemental Table S8). Of the 40,397 predicted transcripts, 29,146 (72.2%) returned a functional annotation. Functional annotation analyses assigned InterPro domain, GO, KEGG pathway, and KEGG ortholog terms to 29,073 (72.0%), 20,580 (50.9%), 8,999 (22.3%) and 7,142 (17.7%) of the predicted transcripts, respectively (Supplemental Table S9).

### Potential candidate genes for primocane-fruiting in blackberry

In blackberry, the primocane-fruiting trait is caused by a single recessive locus that has been mapped between markers FF683693.1 RH_MEa0007aG06 and FF683518.1 RH_MEa0006aC04 in a simple sequence repeat (SSR)-based linkage map of the tetraploid population ‘Prime-Jim^®^’ x ‘Arapaho’ [42]. While these markers were originally placed on linkage group 7 of blackberry, it was later shown that the flanking markers and most others from linkage group 7 of the ‘Prime-Jim^®^’ x ‘Arapaho’ aligned to chromosome 2 of *R. occidentalis* [23]. Based on our genomic data, these markers are located on *R. argutus* chromosome Ra02 at 25,901,374 to 25,901,083 bp (FF683518.1 RH_MEa0006aC04) and 37,085,586 to 37,085,204 bp (FF683693.1 RH_MEa0007aG06). Interestingly, different loci have been found to control primocane-fruiting in raspberry [70] and everbearing flowering in diploid and octoploid strawberries [71, 72], suggesting that flowering in first-year shoots has evolved multiple times in the Rosaceae.

To explore candidate genes for the primocane-fruiting trait in blackberry, blackberry homologs of the *Arabidopsis* flowering time genes listed in FLOR-ID database were mined from the ‘Hillquist’ genome sequence (Supplemental Table S10, [73]). Based on blackberry gene annotations and BLAST analyses, 18 flowering gene homologs were identified within the ∼11.2 Mb primocane-fruiting locus on ‘Hillquist’ chromosome Ra02 (Table 4). Almost half of the genes were involved in epigenetic processes that control gene expression through histone methylation, histone ubiquitinylation, small RNA processing, or as a component of nucleosome assembly. Moreover, six putative transcription factors and three photoperiodic flowering pathway genes (*LATE*, *PRR7*, *CIB4*) were identified in the primocane-fruiting locus.

**Table 4:**
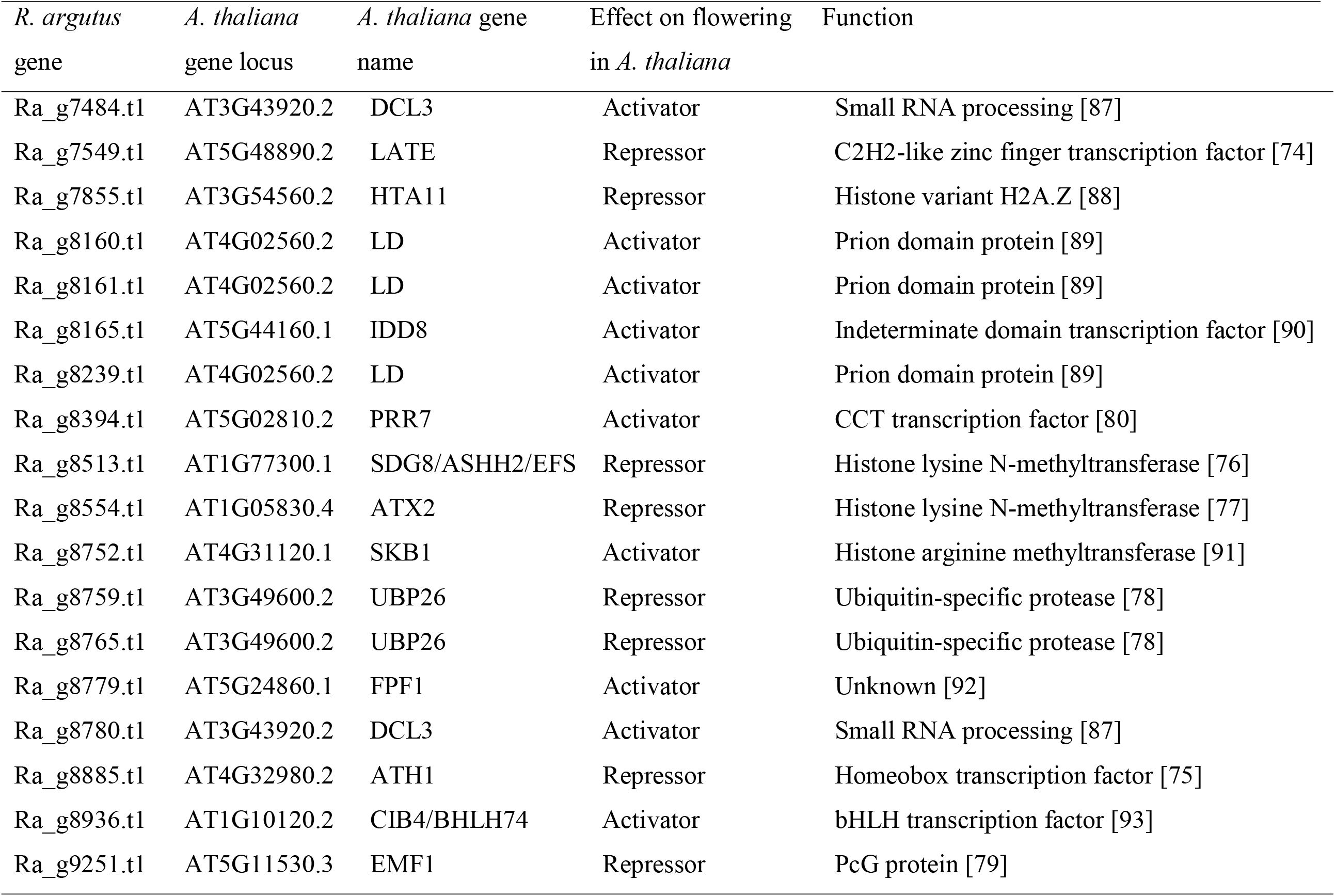
*R. argutus* flowering gene homologs identified in the primocane-fruiting locus from 25.9 to 37.1 Mb on chromosome Ra02.

| *R. argutus*<br>gene | *A. thaliana*<br>gene locus | *A. thaliana* gene<br>name | Effect on flowering<br>in *A. thaliana* | Function |
| --- | --- | --- | --- | --- |
| Ra_g7484.t1 | AT3G43920.2 | DCL3 | Activator | Small RNA processing [87] |
| Ra_g7549.t1 | AT5G48890.2 | LATE | Repressor | C2H2-like zinc finger transcription factor [74] |
| Ra_g7855.t1 | AT3G54560.2 | HTA11 | Repressor | Histone variant H2A.Z [88] |
| Ra_g8160.t1 | AT4G02560.2 | LD | Activator | Prion domain protein [89] |
| Ra_g8161.t1 | AT4G02560.2 | LD | Activator | Prion domain protein [89] |
| Ra_g8165.t1 | AT5G44160.1 | IDD8 | Activator | Indeterminate domain transcription factor [90] |
| Ra_g8239.t1 | AT4G02560.2 | LD | Activator | Prion domain protein [89] |
| Ra_g8394.t1 | AT5G02810.2 | PRR7 | Activator | CCT transcription factor [80] |
| Ra_g8513.t1 | AT1G77300.1 | SDG8/ASHH2/EFS | Repressor | Histone lysine N-methyltransferase [76] |
| Ra_g8554.t1 | AT1G05830.4 | ATX2 | Repressor | Histone lysine N-methyltransferase [77] |
| Ra_g8752.t1 | AT4G31120.1 | SKB1 | Activator | Histone arginine methyltransferase [91] |
| Ra_g8759.t1 | AT3G49600.2 | UBP26 | Repressor | Ubiquitin-specific protease [78] |
| Ra_g8765.t1 | AT3G49600.2 | UBP26 | Repressor | Ubiquitin-specific protease [78] |
| Ra_g8779.t1 | AT5G24860.1 | FPF1 | Activator | Unknown [92] |
| Ra_g8780.t1 | AT3G43920.2 | DCL3 | Activator | Small RNA processing [87] |
| Ra_g8885.t1 | AT4G32980.2 | ATH1 | Repressor | Homeobox transcription factor [75] |
| Ra_g8936.t1 | AT1G10120.2 | CIB4/BHLH74 | Activator | bHLH transcription factor [93] |
| Ra_g9251.t1 | AT5G11530.3 | EMF1 | Repressor | PcG protein [79] |

Ten and eight of the 18 flowering genes in the locus encoded activators and repressors of flowering in *Arabidopsis*, respectively. Floral repressors are primary candidates for primocane-fruiting because a loss-of-function mutation in a repressor could cause this trait to be recessively inherited. Among transcription factors in the locus that repress flowering, LATE is a C2H2 zinc-finger protein that represses the expression of photoperiodic pathway genes *CO* and *FT* [74] and ATH1 is involved in the activation of *FLC* in *Arabidopsis* [75]. Furthermore, many of the identified epigenetic regulators, including STG8, ATX2, UBP26, and EMF1, functioned as floral repressors in *Arabidopsis* by activating the expression of *FLC* [76–79]. No clear *FLC* ortholog was found in the ‘Hillquist’ genome assembly, but these epigenetic regulators likely regulate other targets in blackberry as observed in *Arabidopsis* [77, 79].

Other promising candidate genes identified were *PRR7* and *FD*. *PRR7* encodes a floral activator in *Arabidopsis* [80], and a homologous gene called *BTC1* is involved in the annual to biennial transition in sugar beet [81]. However, if PRR7 controls primocane-fruiting in blackberry, the mechanism is different from beet. In beet, recessive *btc1* alleles confer an obligatory vernalization response and postpone floral initiation into the spring of the second year [81], while recessive alleles of the primocane-fruiting locus cause flowering during the first year in blackberry [82].

Previous studies have shown that *TFL1* encodes a strong repressor of flowering in several Rosaceous species. For example, in diploid woodland strawberry, non-functional *TFL1* alleles cause rapid and perpetual flowering in LD conditions [71]. Similarly, RNA-silencing of *TFL1* orthologs in cultivated strawberry, apple, and pear caused comparative phenotypes in these species [83–85]. Therefore, *TFL1* is also expected to play an important role in the control of flowering in blackberry, and it is a potential target of identified candidate genes. A gene encoding the bZIP transcription factor *FD* was identified just outside the primocane-fruiting locus in the ‘Hillquist’ genome. Recent results show that TFL1 competes for binding to FD with floral activator FT to control common target genes in Arabidopsis [86]. Therefore, a mutation in FD could potentially prevent *TFL1* from repressing floral activators that are needed for floral initiation during the first season in primocane-fruiting genotypes, leading to the observed phenotype.

## Conclusions

The first high-quality chromosome-length genome assembly and annotation of the diploid blackberry *R. argutus* ‘Hillquist’ is reported in this manuscript. Comparisons of the ‘Hillquist’ genome with the related species *R. idaeus* [26] and *R. chingii* [27] demonstrated that the Hi-C assembly represented the majority of the genome and was of high quality. BUSCO analysis and comparisons of predicted genes with other *Rubus* genomes showed that the structural and functional annotations of the assembly were also comprehensive. Analysis of repeat content revealed that approximately 52.8% of the genome was composed of repetitive elements and that the *Gypsy* superfamily of LTR-REs accounted for the largest fractions of the genome. Developing new GBS-based maternal haplotype map of the tetraploid blackberry breeding selection A-2551TN that was highly collinear with the physical sequence of ‘Hillquist’ demonstrated the utility of this new genome for molecular breeding applications in tetraploid fresh-market blackberries. The new ‘Hillquist’ genome assembly and its annotation were also used to identify potential candidate genes for the economically important trait of primocane-fruiting. The ‘Hillquist’ genome sequence and annotation presented here will assist blackberry breeders and scientists in marker development and genomic-assisted breeding and facilitate future studies of *Rubus* biology, genetics, and genomics.

## Availability of Supporting Data and Materials

Raw sequencing data and the genome assembly of *R. argutus* presented here are available at the NCBI under Bioproject ID PRJNA830911. Hi-C data are available on Bioproject PRJNA512907 (SRX8934844). Interactive Hi-C contact maps of the ‘Hillquist’ genome sequence assembly are available via the www.dnazoo.org website (https://www.dnazoo.org/assemblies/Rubus_argutus). The ‘Hillquist’ genome assembly and annotation can also be accessed at the Genome Database for Rosaceae (https://www.rosaceae.org/Analysis/13328362; [39]) under the accession number tfGDR1055.

## Abbreviations

BLAST: Basic Local Alignment Search Tool
bp: base pairs
BUSCO: Benchmarking Universal Single-Copy Orthologs
BWA: Burrows-Wheeler Aligner
CCS: circular consensus sequence
cDNA: complementary DNA
CTAB: cetyl trimethylammonium bromide
EMBOSS: European Molecular Biology Open Software Suite
Gb: gigabase pairs
GBS: genotyping-by-sequencing
GC: guanine-cytosine
GDR: Genome Database for Rosaceae
GO: gene ontology
Hi-C: high-throughput chromosome conformation capture
kb: kilobase pairs
lORF: longest open reading frame
LTR: long terminal repeat
Mb: megabase pairs
MYA: million years ago
NCBI: National Center for Biotechnology Information
NCGR: National Clonal Germplasm Repository
NCSU: North Carolina State University
ORF: open reading frame
PacBio: Pacific Biosciences
RE: retrotransposable element
RNA-Seq: RNA-sequencing
SMRT: single molecule real-time
SNP: single nucleotide polymorphism
SSR: simple sequence repeat
TAIR: The Arabidopsis Information Resource
USDA-ARS: United States Department of Agriculture Agricultural Research Service.

## Authors’ Contributions

T.B. and A.L. performed structural annotation analyses. R.A. performed DNA and RNA extractions, library preparation, PacBio genome assembly, and genome quality assessments. O.D., M.P., D.W. and E.L.A. performed the in situ Hi-C experiment, Hi-C-guided assembly, and associated analyses. M.Bu. performed functional annotation analyses. A.C., F.M., G.U., and L.N performed analyses of repeat content. T.H. and J.A. analyzed potential candidate genes for primocane-fruiting. J.D. performed synteny analysis with other Rosoideae genome assemblies. N.B. coordinated sample collection and flow cytometry. M.W., M.A., and B.O., conducted genotyping-by-sequencing and linkage mapping. T.P. and C.B. conducted IsoSeq analysis. D.M. coordinated and assisted with 10x, Hi-C sequencing, RNA-Seq, and genome assembly. D.J.S. coordinated and assisted with annotation, repeat content, and flowering gene analyses. M.W. conceived the study and M.W., D.J.S., D.M., M.Bo., E.A., N.B., G.F., and H.A. coordinated research and provided conceptual guidance. M.W., T.B., D.J.S., A.C., and T.H. authored the manuscript. All authors approved the final manuscript.

## Acknowledgments

Hi-C data were created by the DNA Zoo Consortium (www.dnazoo.org). DNA Zoo sequencing effort is supported by Illumina, Inc. We acknowledge Kim Hummer and Jill Bushakra from the USDA-ARS, National Clonal Germplasm Repository, who assisted with sample collection and plant maintenance, John Clark and the University of Arkansas System Division of Agriculture Fruit Research Station staff who developed the A-2551TN x APF-259TN mapping population and conducted routine plant maintenance, Felicidad Fernández-Fernández and Mario Caccamo from NIAB-EMR who participated in fruitful conversation and gave valuable advice at the inception of this project, and Ryan Rapp, Cherie Ochsenfeld, Aabid Shariff, Gina Pham and Xiaoyu Zhang from Pairwise for their assistance with IsoSeq analyses. We thank Mike Stratton and Julia Wilson for their continuing support for the 25 genomes for 25 years project. We also thank Michelle Smith, Craig Corton, and Karen Oliver for their work in processing the samples for 10X and RNA-seq.

## Additional Files

Supplemental File S1. Full methods and results for development of the A-2551TN maternal haplotype map.

Supplemental Table S1. Summary of sequencing data from PacBio, Hi-C, and 10X platforms.

Supplemental Table S2. Summary statistics for the seven chromosome-length scaffolds corresponding to the ‘Hillquist’ blackberry (*R. argutus*) base chromosomes.

Supplemental Table S3. Depth of genotyping-by-sequencing read coverage in the parents and F_1_ progeny of the A-2551TN x APF-259TN mapping population and percent of reads aligning to unique positions in the ‘Hillquist’ blackberry (*R. argutus*) and black raspberry (*R. occidentalis*) reference genomes.

Supplemental Table S4. Distribution of single-dose allele markers across the A-2551TN maternal haplotype map.

Supplemental Table S5. Marker positions in the A-2551TN maternal haplotype map, physical positions on the ‘Hillquist’ blackberry (*R. argutus*) genome assembly, and genotype scores in the parents and progeny.

Supplemental Table S6. Accession numbers corresponding to Sequence Reads Archive (SRA) and assembly data of the four Rosaceae species used for comparative analysis of repeat composition with the ‘Hillquist’ blackberry (*R. argutus*) genome assembly. *Potentilla micrantha* datasets were retrieved from the GigaScience GigaDB repository.

Supplemental Table S7. Number and classification of full-length long terminal repeat retrotransposons (LTR-REs) identified in the genomes of five Rosaceae species.

Supplemental Table S8. Predicted proteins in the *R. argutus* genome assembly along with physical positions and matches obtained from blastp analyses with nr, Araport11, RefSeq, SwissProt and TrEMBL databases as subjects.

Supplemental Table S9. Summary of functional annotation of ‘Hillquist’ gene predictions including transcripts with matches to InterPro domains, Gene Ontology (GO) terms, KEGG pathway and KEGG orthologs.

Supplemental Table S10. *R. argutus* blackberry homologs of *Arabidopsis* flowering time genes listed in FLOR-ID database.

Supplemental Figure S1. GenomeScope k-mer analysis with 10x Genomics Illumina paired-end reads.

Supplemental Figure S2. Whole-genome alignment plot between the ‘Hillquist’ blackberry (*R. argutus*) genome assembly and the chromosome-scale assembly of *R. chingii*.

Supplemental Figure S3. Pedigree of the A-2551TN x APF-259TN mapping population.

Supplemental Figure S4. The 30 linkage groups of the A-2551TN maternal haplotype map. Marker positions are expressed in cM.

Supplemental Figure S5. The amount of sequence repeat-masked by repeats grouped by their length.

Supplemental Figure S6. Flowchart of the structural gene annotation.

Supplemental Figure S7. Distribution of the number of multiple alternative isoforms per protein-coding locus. There are 36,836 genes without alternative isoforms.

Supplemental Figure S8. BUSCO annotation assessment of the ‘Hillquist’ blackberry (*R. argutus*) genome assembly.

