## Supplemental File S1 for "A chromosome-length genome assembly and annotation of blackberry (*Rubus argutus*, cv. ‘Hillquist’)"

**Supplemental File 1: Full methods and results for the development of the A-2551TN maternal haplotype map**

*Plant material.* The A-2551TN x APF-259TN mapping population for this study was grown at the University of Arkansas System Division of Agriculture (UA) Fruit Research Station, Clarksville [west-central Arkansas, lat. 35º31'5"N, long. 93º24'12"W; U.S. Department of Agriculture (USDA) plant hardiness zone 7b (USDA, 2012). The two parents were crossed in 2015 to create 164 $F_1$ individuals. The cross was repeated in 2016 to create 86 $F_1$ individuals, for a total population of 250 progeny. Both parents were advanced breeding selections from the UA Fruit Breeding Program selected for their short stature and reduced internode length for ornamental applications. Inbreeding coefficients were calculated for A-2551TN, APF-259TN, and their progeny based on pedigree records using breedr (Supplemental Figure S3; https://github.com/mchizk1/breedr).

*Genotyping-by-sequencing.* DNA was extracted from young leaf samples harvested from parents and progeny following a modified CTAB protocol [1]. The extractions were quantified by a Qubit® fluorometer (Thermo Fisher Scientific, Waltham, MA, USA) and diluted to a concentration of 200 ng/µL in 30 µL wells. A modified genotyping-by-sequencing (GBSpoly) library preparation and sequencing was performed at the Genomic Sciences Laboratory, North Carolina State University (Raleigh, NC). The GBSpoly library preparation protocol is optimized for polyploids and highly heterozygous genomes. It is a quantitative assay that produces uniform coverage across samples and loci, and hence, allows for allele dosage estimation based on ratios of allelic composition and avoids minimal missing rate typically associated similar protocols [2,3]. Briefly, the frequent and rare cutter restriction enzymes, *Cvi*AII and *Tse*I, respectively, were used to digest the DNA samples. The digested DNA samples were purified with AMPure®

XP magnetic beads (Beckman Coulter Inc., Brea, CA) and the resulting fragments were ligated to barcoded adapters, with 6 bp buffer sequences upstream of the barcodes, which ranged from 6-9 bp. The buffer sequences were included to decrease the base call error rate in the barcode region and minimize barcode swapping (i.e. avoids misassigning reads to individuals) during demultiplexing. A post-ligation digest with *Cvi*AII/*Tse*I was then performed to eliminate chimeric sequences. Following the second digest, the pooled libraries were purified with AMPure® XP magnetic beads and selected for 300-400 bp fragments using Pippin Prep (Sage Sciences Inc., Beverly, MA) to minimize PCR fragment length bias and limit the proportion of the genome that is captured for genotyping (higher density can be achieved by increasing fragment size window selection).

The 250 progeny and parents were first pooled in groups of 48 samples. Each pool was sequenced on a HiSeq 2500 (Illumina, San Diego, CA) lane with parents sequenced at 8x higher coverage than the progeny to ensure accurate parental dosage calls could be made for all polymorphic SNPs. Sequencing read depth was uneven across samples and inadequate for many genotypes in this first sequencing run. Thus, 188 progeny and parents (2x) were pooled in groups of 96 samples and sequenced on two NovaSeq™ 6000 System (Illumina, San Diego, CA) lanes.

*Genotype calling*. Raw Fastq files were preprocessed using ngsComposer (https://github.com/bodeolukolu/ngsComposer [4]). Preprocessing steps included buffer sequence trimming, demultiplexing, filtering based on presence of restriction site motif, end trimming, adapter removal, and quality threshold filtering. The GBSapp pipeline was then used for SNP-calling and filtering (https://github.com/bodeolukolu/GBSapp; [2]). Processed reads were aligned to black raspberry (*R. occidentalis* L.) [5] and 'Hillquist' genomes using BWA-MEM [6]. Alignment files were then pre-processed before variant calling with GATK

HaplotypeCaller [7] using the GVCF mode for single sample calling, and then followed by downstream joint genotyping using the GATK GenomicsDBImport and GenotypeGVCFs tools. Genotypes with less than 25 reads for each variant were called as missing because a greater number of reads are required to make accurate genotypic calls in tetrasomic polyploids than diploid populations. Markers (SNPs and InDels) and genotypes with greater than 20% missing data were initially removed as well as markers that deviated from expected segregation ratios at $P < 0.001$.

*Pseudo-testcross mapping*. A maternal haplotype map of A-2551TN was created in JoinMap 4.1 following the pseudo-testcross strategy [8,9]. Only markers that were heterozygous in the simplex condition (1/0/0/0) in A-2551TN and homozygous in the nulliplex condition (0/0/0/0) in APF-259TN were used to construct the haplotype-resolved maternal linkage map. Prior to map construction, individuals with 20% or more of missing data in the mapping dataset were excluded. Individuals with ratios of homozygote to heterozygote calls greater than 2:1 or less than 1:2 in the mapping dataset were identified as possible selfed progeny of A-2551TN or accidental outcrosses with contaminant pollen from other sources and removed from the mapping dataset. Identical markers and markers that deviated from expected segregation ratios according to the $\chi^2$ test ($P < 0.10$) were excluded from mapping. JoinMap 4.1 can only handle datasets of 4,000 or fewer markers. Because the number of markers that passed initial missing data and segregation distortion thresholds for the A-2551TN map far exceeded 4,000, 5% was set as the maximum allowable missing data for each marker.

The threshold linkage logarithm of odds (LOD) for establishing initial groups was set to 9.0. Marker order and distances were then determined using the regression mapping algorithm with default settings and Haldane’s mapping function. There was insufficient linkage in the data to create maps for several of the linkage groups that clustered together at LOD 9.0 in the A-2551TN haplotype map. In these instances, higher LODs (ranging from 10-17) were selected for establishing groups with sufficient linkages for mapping. The JoinMap 4.1 regression mapping algorithm can only be used to order linkage groups of up to 250 markers. Therefore, in instances where more than 250 markers were assigned to a linkage group, markers with greater than 95% similarity were excluded from mapping. Charts of the genetic linkage map were drawn using MapChart 2.1 [10]. Plots aligning the maternal haplotype map to the 'Hillquist' reference genome were generated in the R package *ggplot2* [11].

**Results**

*Genotype calling*. Post-filtering, a total of 615.4 million sequencing reads were obtained for the parents and progeny from both the HiSeq 2500 and NovaSeq™ 6000 System sequencing runs. After demultiplexing, processing, and quality filtering, we obtained 8 million reads for A-2551TN, 8.7 million reads for APF-259TN, and an average of 2.1 million reads for each of the progeny. On average, 85.9% of reads were mapped to unique positions on the 'Hillquist' genome and 67.3% of reads mapped to unique positions on the black raspberry genome (Supplemental Table S3). 1,811,617 and 2,022,664 polymorphic markers were identified when these reads were aligned to the black raspberry and 'Hillquist' genomes, respectively, using the GBSapp pipeline. Only the markers identified using the 'Hillquist' reference genome were used for mapping. Two hundred and two of the original 250 progeny and 14,492 markers passed the initial filters for missing data and segregation distortion. Of these markers, 8,699 (58%) were classified as single-dose markers segregating in A-2551TN, 2,003 (13%) were classified as single-dose markers segregating in APF-259TN, 2,198 (15%) were classified as double-simplex (single-dose markers segregating in both parents), and 2,092 (14%) were classified as multiplex.

*Genetic linkage maps*. Only 119 of the original 250 progeny remained in the mapping dataset after filtering for greater than 20% missing single-dose markers, for ratios of homozygote to heterozygote calls greater than 2:1 or less than 1:2, and for progenies that are potential selfs. Because JoinMap 4.1 can only handle datasets with fewer than 4,000 markers. Of the 8,699 single-dose markers segregating in A-2551TN passed initial quality filtering, we excluded all markers with greater than 5% missing data for the maternal map. Of the 3,796 markers used for linkage mapping in the maternal haplotype map, 470 were removed because they were identical, 201 were ungrouped, and 3,125 were placed in 30 linkage groups. Originally 395 markers were placed in linkage group 6a and 219 markers in linkage group 6b, but the regression mapping algorithm in JoinMap 4.1 could not process the ordering of over 250 markers per groups so markers with over 95% similarities were removed. Therefore, the final maternal haplotype map was composed of 2,935 markers, with between 5 and 249 markers per linkage group (Supplemental Tables S4, S5; Supplemental Figure S4). The total map length was 2,411.81 cM with linkage groups ranging from 18.61 cM to 146.65 cM in length and an average of 1 marker every 0.82 cM.

The physical positions of the mapped markers in the 'Hillquist' reference genome were used to identify homologous linkage groups for each of the seven base chromosomes of blackberry. In general, the genetic and physical maps were strongly collinear, with no major translocations or inversions (Figure 4). Four homologous linkage groups were found as expected for chromosomes 1, 2, 3, 4, and 6 in the maternal haplotype map, but five homologous linkage groups corresponding to chromosomes 5 and 7 were identified. While many of the linkage groups in the A-2551TN maternal haplotype map contained markers that aligned to physical positions across the length of the chromosome, 10 linkage groups had markers aligned to physical positions spanning less than 10 Mbp in the 'Hillquist' genome. Based on the physical positions of these markers on short linkage groups, it is likely that linkage groups 7b and 7d and linkage groups 5c and 5e actually belong to the same haplotype of A-2551TN. Inbreeding coefficients for A-2551TN, APF-259TN, and their progeny were 0.101, 0.132, and 0.099, respectively. The relatively high inbreeding coefficients in both parents and the progeny were the most likely cause of the short linkage groups and gaps in the haplotype maps.

A high degree of collinearity between the diploid 'Hillquist' reference and the tetraploid mapping population was expected considering 'Hillquist' is highly represented in the pedigrees of both parents of the mapping populations (Supplemental Figure S3). Still, the agreement between the physical map of 'Hillquist' and the tetraploid haplotype map developed in this study validates the order and orientation of the HiC-based chromosome scale assembly of 'Hillquist' and its utility for genomic breeding research in polyploid fresh-market blackberries.
