## Supplemental Table S1 for "A chromosome-length genome assembly and annotation of blackberry (*Rubus argutus*, cv. ‘Hillquist’)"

Supplemental Table S1. Summary of PacBio sequencing data

| | PacBio |
|---|---|
| Number of Bases | 25,862,352,946 |
| Number of Reads | 3,789,829 |
| N50 Read Length (bp) | 12,250 |
| Mean Read Length (bp) | 6,824 |
| GC % | 38% |
| Nominal coverage (337 Mb genome) | 77x |
