## Supplemental Table S2 for "A chromosome-length genome assembly and annotation of blackberry (*Rubus argutus*, cv. ‘Hillquist’)"

Supplemental Table S2. Summary statistics for the seven super-scaffolds corresponding to the 'Hillquist' blackberry (*R. argutus*) base chromosomes.

| Chromosomes | Total Length (bp) | N count | Gaps |
|---|---|---|---|
| Ra01 | 31951077 | 48000 | 96 |
| Ra02 | 39129706 | 57500 | 115 |
| Ra03 | 41973286 | 54000 | 108 |
| Ra04 | 36116165 | 51000 | 102 |
| Ra05 | 38624057 | 41500 | 83 |
| Ra06 | 45459684 | 54000 | 108 |
| Ra07 | 37018114 | 51500 | 103 |
| Total size (7 chromosomes) | 270272089 | 357500 | 715 |
| Unassembled fragments (343 scaffolds) | 27965006 | 20000 | 40 |
| Total genome | 298237095 | 377500 | 755 |
