## Supplemental Table S4 for "A chromosome-length genome assembly and annotation of blackberry (*Rubus argutus*, cv. ‘Hillquist’)"

Supplemental Table S4. Distribution of single-dose allele markers across the A-2551TN maternal haplotype map.

| Linkage group[z] | No. markers[y] | Length (cM) | Physical positions of mapped markers (Mb) |
|---|---|---|---|
| 1a | 169 | 127.31 | 0.19-31.38 |
| 1b | 58 | 76.42 | 0.96-46.67 |
| 1c | 32 | 36.15 | 16.30-20.38 |
| 1d | 24 | 18.61 | 3.54-4.82 |
| 2a | 195 | 113.66 | 0.24-31.98 |
| 2b | 100 | 65.26 | 1.89-36.99 |
| 2c | 26 | 74.48 | 6.70-28.55 |
| 2d | 14 | 61.52 | 0.23-26.53 |
| 3a | 123 | 125.99 | 4.19-41.83 |
| 3b | 120 | 146.65 | 2.06-41.46 |
| 3c | 9 | 18.71 | 10.18-11.74 |
| 3d | 5 | 28.60 | 16.22-19.49 |
| 4a | 147 | 65.81 | 23.54-34.61 |
| 4b | 119 | 72.76 | 13.49-32.96 |
| 4c | 79 | 91.95 | 0.90-31.00 |
| 4d | 48 | 44.16 | 22.37-27.44 |
| 5a | 170 | 97.21 | 1.64-34.58 |
| 5b | 151 | 130.18 | 0.10-38.53 |
| 5c | 89 | 27.73 | 0.98-3.57 |
| 5d | 54 | 91.67 | 6.12-35.11 |
| 5e | 32 | 43.87 | 29.96-37.62 |
| 6a | 249 | 88.74 | 17.27-45.25 |
| 6b | 175 | 144.62 | 0.32-45.43 |
| 6c | 165 | 84.35 | 3.08-42.41 |
| 6d | 102 | 88.86 | 9.16-45.25 |
| 7a | 219 | 132.84 | 0.13-36.48 |
| 7b | 134 | 70.18 | 0.09-25.41 |
| 7d | 35 | 48.33 | 29.49-32.06 |
| 7e | 25 | 106.83 | 0.38-36.94 |
| Total | 2935 | 2411.81 | - |

[z] Linkage LOD 9.0 was used to establish baseline linkage groups. Higher LOD thresholds were imposed in three cases where there was insufficient linkage in the data to create maps for linkage groups that clustered together at LOD 9.0 in the A-2551TN maternal haplotype map, including 5b and 5d (split at LOD 10), 6c and 6d (split at LOD 17), and 7b, 7c, and 7d (split at LOD 11).

[y] Originally 395 markers were placed in linkage group 6a and 219 markers in linkage group 6b in the maternal haplotype map, but markers with over 95% similarities were removed because the regression mapping algorithm in JoinMap 4.1 could not handle ordering over 250 markers per group.
