## Supplemental Table S6 for "A chromosome-length genome assembly and annotation of blackberry (*Rubus argutus*, cv. ‘Hillquist’)"

Supplemental Table S6. Accession numbers corresponding to assembly data of the four Rosaceae species used for comparative analysis of repeat composition with the 'Hillquist' blackberry (*R. argutus*) genome assembly. *Potentilla micrantha* datasets were retrieved from the GigaScience GigaDB repository.

| Species | Assembly |
|---|---|
| *Fragaria vesca* | GCF_000184155.1 |
| *Potentilla micrantha* | http://gigadb.org/dataset/100407 |
| *Prunus persica* | GCF_000346465.2 |
| *Malus domestica* | NC_041798.1 |
