## Supplemental Table S7 for "A chromosome-length genome assembly and annotation of blackberry (*Rubus argutus*, cv. ‘Hillquist’)"

Supplemental Table S7. Number and classification of full-length long terminal repeat retrotransposons (LTR-REs) identified in the genomes of five Rosaceae species.

| **CLASS I - LTR-RE** | ***Rubus argutus*** | ***Fragaria vesca*** | ***Potentilla micrantha*** | ***Prunus persica*** | ***Malus domestica*** |
|---|---|---|---|---|---|
| ***Gypsy*** | **217** | **54** | **233** | **124** | **1,559** |
| *chromovirus\|CRM* | 6 | 16 | 3 | 20 | 4 |
| *chromovirus\|Galadriel* | 2 | 2 | 1 | ND | ND |
| *chromovirus\|Reina* | 21 | 2 | 16 | 4 | 60 |
| *chromovirus\|Tekay* | 12 | 2 | 4 | 13 | 58 |
| *non-chromovirus\|OTA\|Athila* | 102 | 1 | 7 | 4 | 187 |
| *non-chromovirus\|OTA\|Ogre/Tat* | 74 | 31 | 202 | 83 | 1250 |
| ***Copia*** | **409** | **144** | **228** | **564** | **1,097** |
| *Ale* | 30 | 19 | 18 | 171 | 197 |
| *Alesia* | 1 | ND | ND | ND | 25 |
| *Angela* | 2 | ND | ND | ND | 154 |
| *Bianca* | 274 | 102 | 157 | 267 | 346 |
| *Ikeros* | 39 | 2 | 12 | 2 | 62 |
| *Ivana* | 22 | 13 | 30 | 100 | 191 |
| *SIRE* | 20 | 1 | 6 | 14 | 4 |
| *TAR* | ND | ND | ND | 2 | 5 |
| *Tork* | 21 | 7 | 5 | 8 | 113 |
| **Unclassified** | **10** | **6** | **2** | **6** | **6** |
| **Total LTR-RE** | **636** | **204** | **463** | **694** | **2,662** |
