## Supplemental Table S9 for "A chromosome-length genome assembly and annotation of blackberry (*Rubus argutus*, cv. ‘Hillquist’)"

Supplemental Table S9. Summary of functional annotation of ‘Hillquist’ gene predictions including transcripts with matches to InterPro domains, Gene Ontology (GO) terms, KEGG pathway and KEGG orthologs.

| | Total | All databases | InterPro domain | GO terms | KEGG pathway | KEGG ortholog |
|---|---|---|---|---|---|---|
| Whole complement of predicted transcripts | 40397 | 29146 (72.2%) | 29073 (72.0%) | 20580 (50.9%) | 8999 (22.3%) | 7142 (17.7%) |
| Transcripts predicted *ab initio* & some external support | 31326 | 27012 (86.2%) | 26942 (86.0%) | 19363 (61.8%) | 8930 (28.5%) | 7098 (22.7%) |
| Transcripts predicted *ab initio* & full external support | 19937 | 17826 (89.4%) | 17781 (89.2%) | 12918 (64.8%) | 7273 (36.5%) | 5812 (29.2%) |
| Transcripts predicted *ab initio* only | 9407 | 2297 (24.4%) | 2294 (24.4%) | 1313 (14.0%) | 93 (1.0%) | 60 (0.1%) |
