## Supplemental Figures for "A chromosome-length genome assembly and annotation of blackberry (*Rubus argutus*, cv. ‘Hillquist’)"

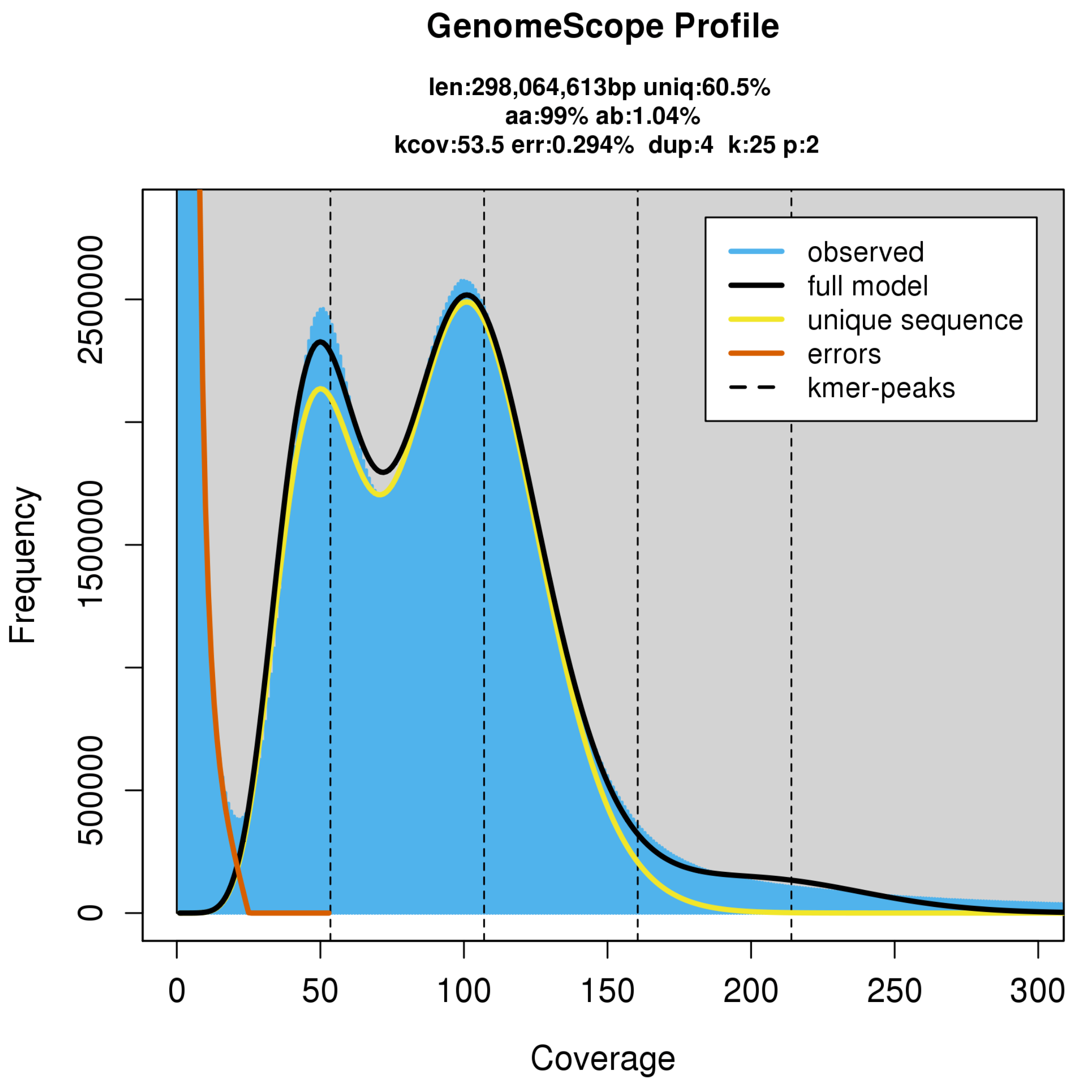


Supplemental Figure S1. GenomeScope k-mer analysis with 10x Genomics reads.

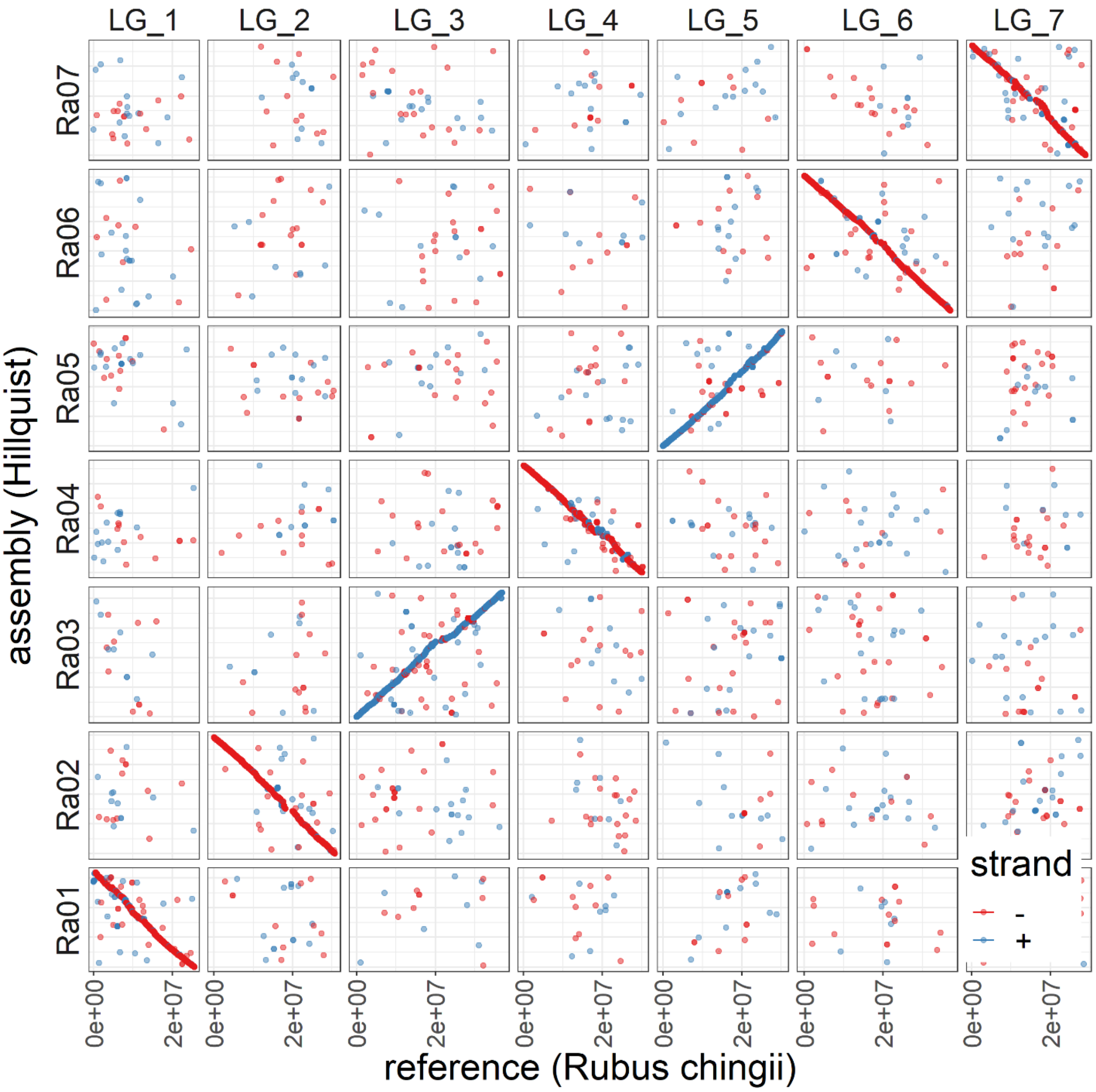


Supplemental Figure S2. Whole-genome alignment plot between the 'Hillquist' blackberry (*R. argutus*) genome assembly and the chromosome-scale assembly of *R. chingii.*

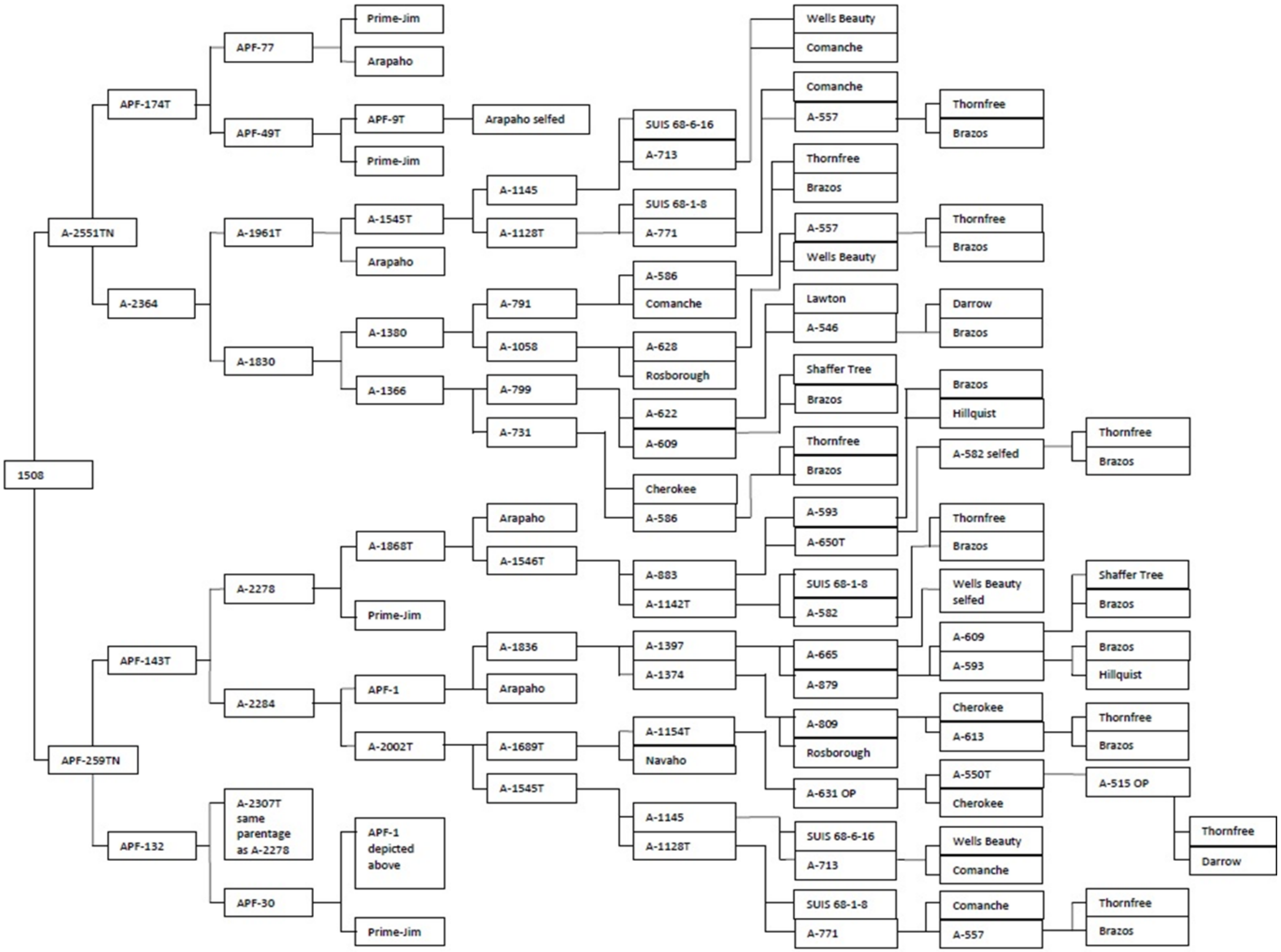


Supplemental Figure S3. Pedigree of the A-2551TN x APF-259TN mapping population.

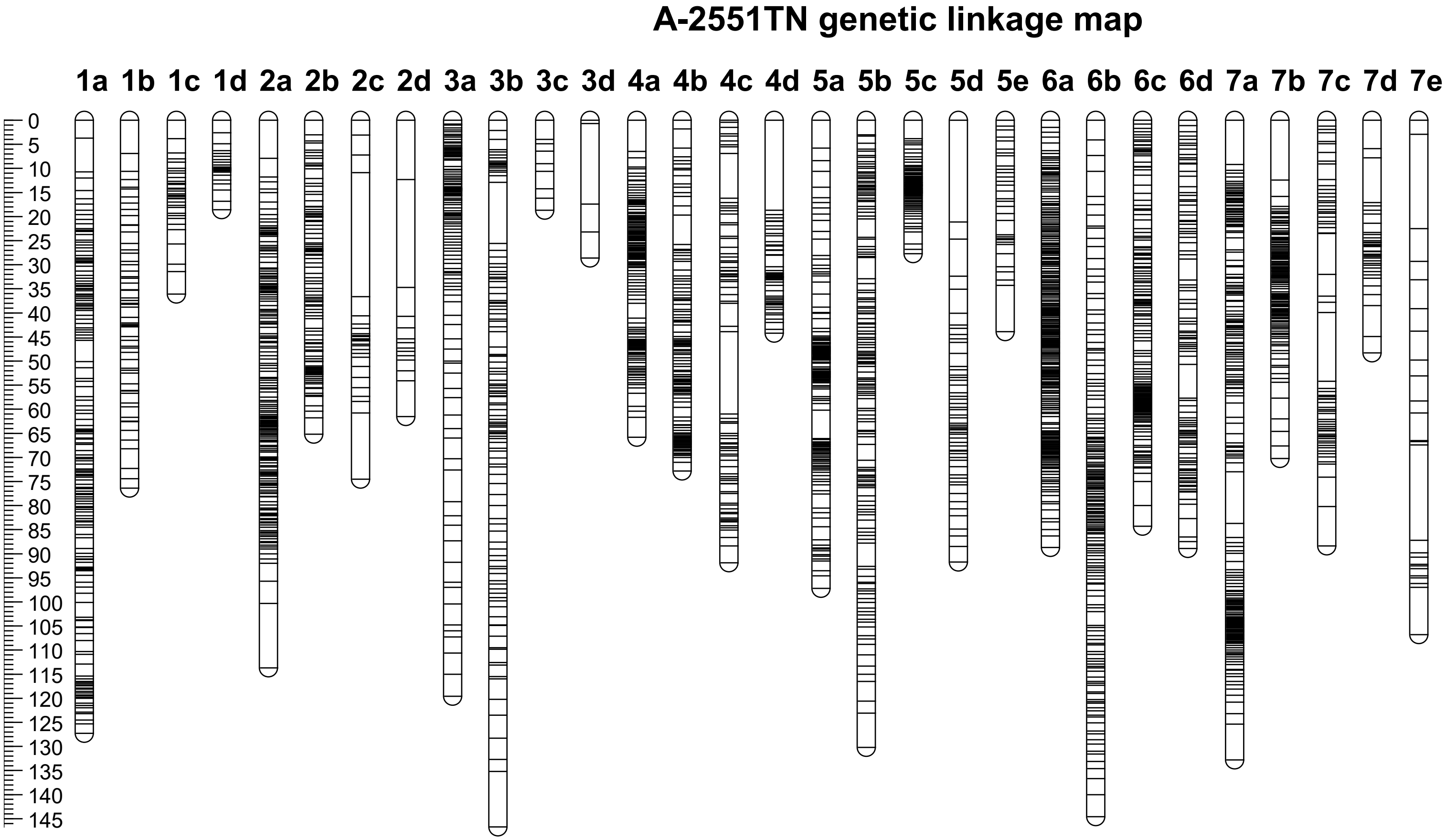


Supplemental Figure S4. The 30 linkage groups of the A-2551TN maternal haplotype map. Marker positions are expressed in cM.

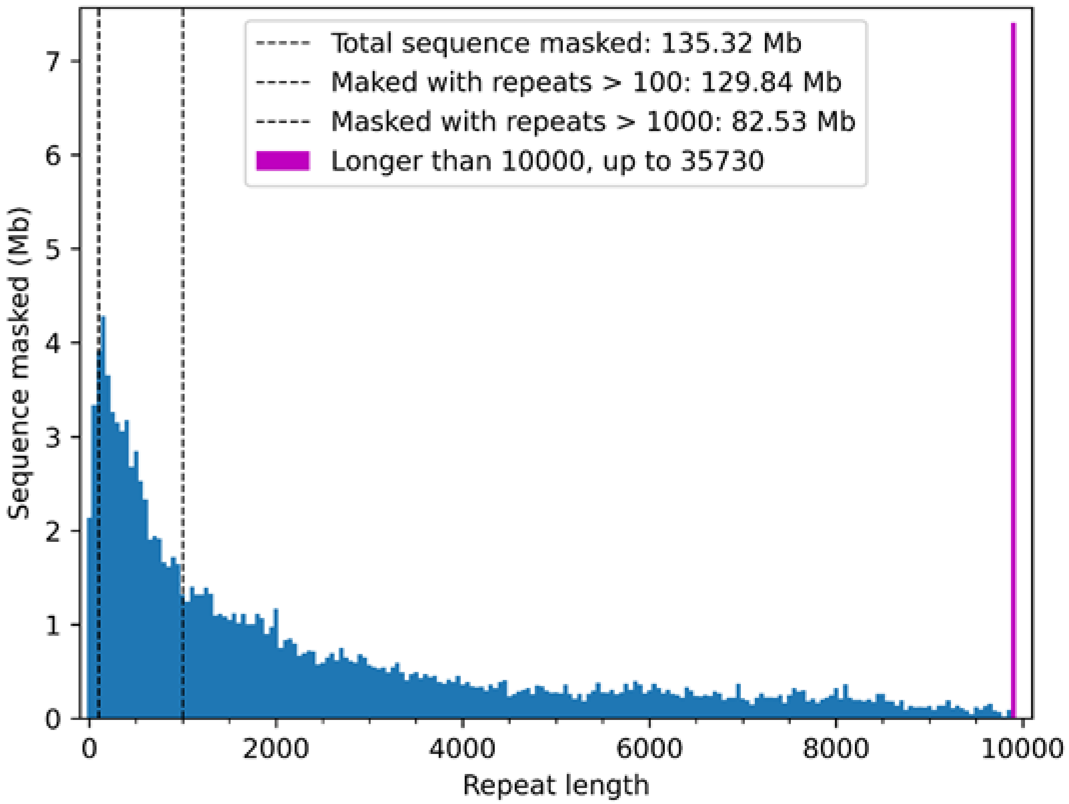


Supplemental Figure S5. The amount of sequence repeat-masked by repeats grouped by their length.

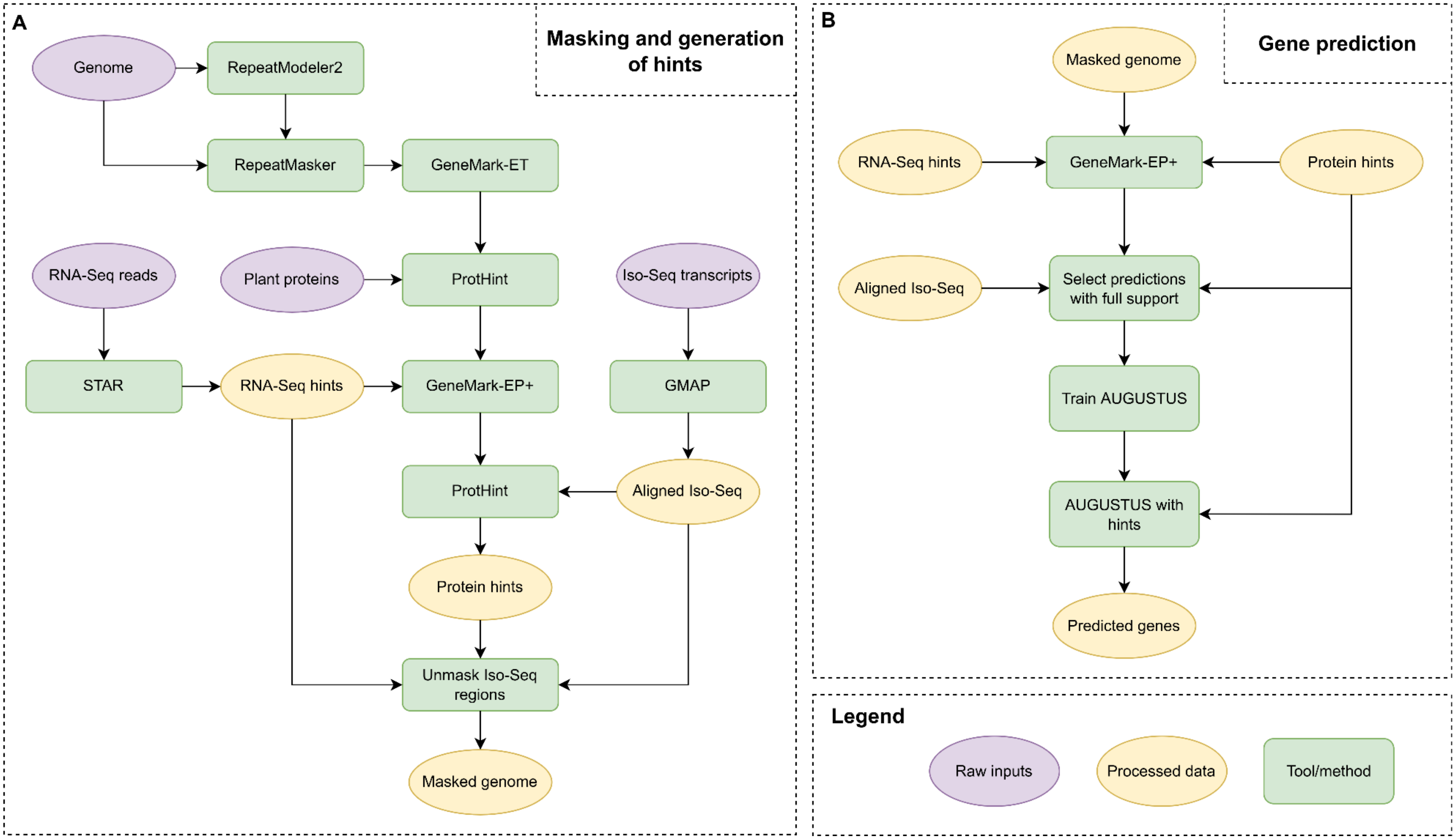


Supplemental Figure S6. Flowchart of the structural gene annotation.

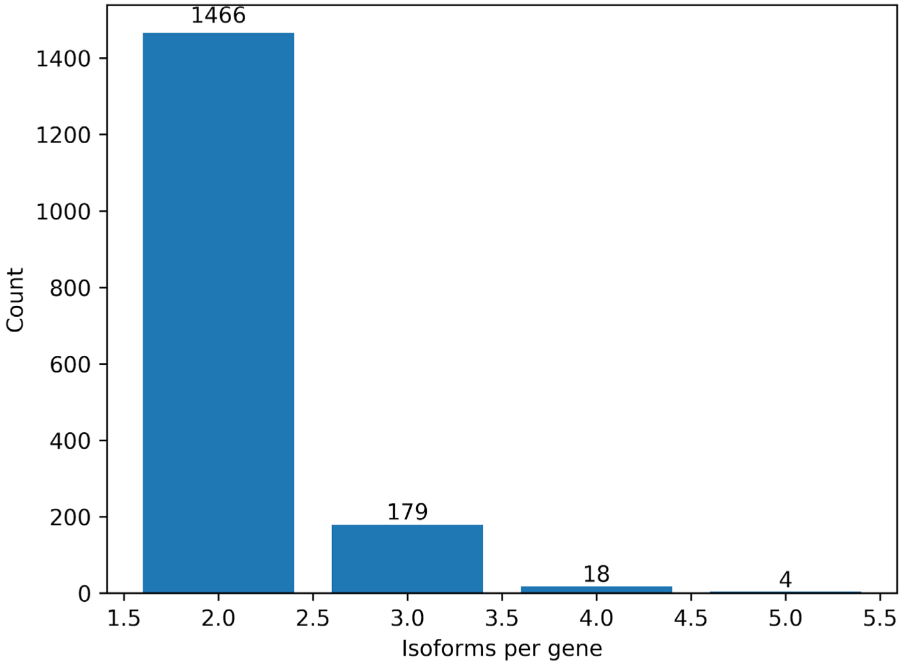


Supplemental Figure S7. Distribution of the number of multiple alternative isoforms per protein-coding locus. There are 36,836 genes without alternative isoforms.

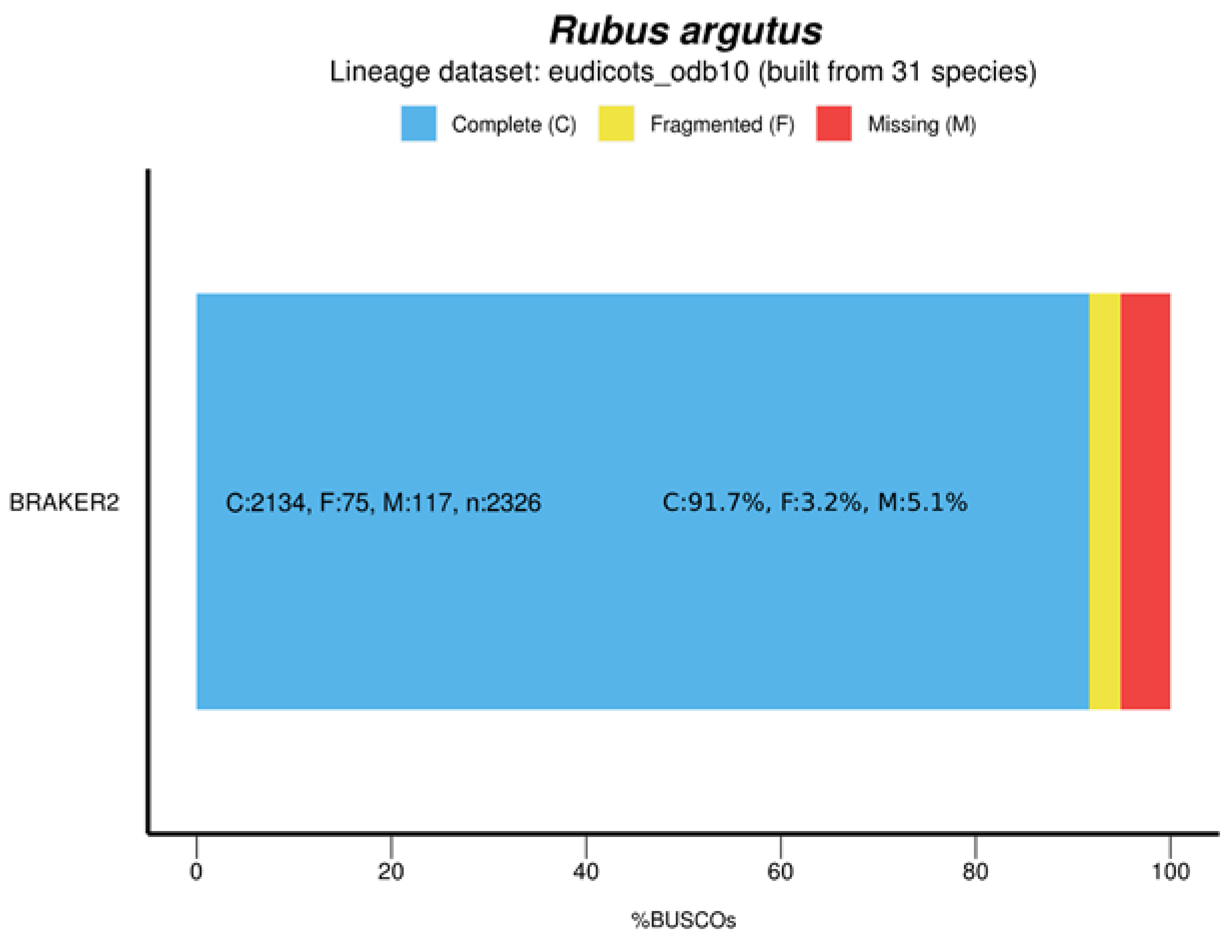


Supplemental Figure S8. BUSCO annotation assessment.
